# Genomic and single-cell characterization of patient-derived tumor organoid models of head and neck squamous cell carcinoma

**DOI:** 10.1101/2024.06.28.601068

**Authors:** Jung Hyun Um, Yueyuan Zheng, Qiong Mao, Chehyun Nam, Hua Zhao, Yoon Woo Koh, Su-Jin Shin, Young Min Park, De-Chen Lin

## Abstract

Head and Neck Squamous Cell Carcinoma (HNSCC) remains a significant health burden due to tumor heterogeneity and treatment resistance, emphasizing the need for improved biological understanding and tailored therapies. This study enrolled 31 HNSCC patients for the establishment of patient-derived tumor organoids (PDOs), which faithfully maintained genomic features and histopathological traits of primary tumors. Long-term culture preserved key characteristics, affirming PDOs as robust representative models. PDOs demonstrated predictive capability for cisplatin treatment responses, correlating *ex vivo* drug sensitivity with patient outcomes. Bulk and single-cell RNA sequencing unveiled molecular subtypes and intratumor heterogeneity (ITH) in PDOs, paralleling patient tumors. Notably, a hybrid epithelial-mesenchymal transition (hEMT)-like ITH program is associated with cisplatin resistance and poor patient survival. Functional analyses identified amphiregulin (AREG) as a potential regulator of the hybrid epithelial/mesenchymal state. Moreover, AREG contributes to cisplatin resistance via EGFR pathway activation, corroborated by clinical samples. In summary, HNSCC PDOs serve as reliable and versatile models, offer predictive insights into ITH programs and treatment responses, and uncover potential therapeutic targets for personalized medicine.

**One Sentence Summary:** This study establishes patient-derived tumor organoids (PDOs) from 31 Head and Neck Squamous Cell Carcinoma (HNSCC) patients, faithfully recapitulating characteristics of primary tumors and accurately predicting clinical responses to cisplatin treatment. We reveal intertumoral heterogeneity within PDOs and a hybrid epithelial-mesenchymal transition (hEMT) program conferring cisplatin resistance, highlighting amphiregulin (AREG) as a regulator of cellular plasticity and potential therapeutic target for HNSCC treatment.

## Introduction

Head and neck squamous cell carcinoma (HNSCC) is a malignant tumor originating from the squamous cells covering mucous membranes of the oral cavity, pharynx, and larynx; HNSCC accounts for more than 90% of all head and neck cancers (*1*). The incidence of HNSCC varies by region and is associated with tobacco exposure, alcohol consumption, or human papillomavirus infection (HPV) (*2*). Since a large number of HNSCC patients are diagnosed in an advanced stage, they require multi-modality treatments that integrate surgery, chemotherapy, and radiation. Cisplatin is a standard drug used for either definitive or adjuvant concurrent chemoradiotherapy (CCRTx) and it can increase the therapeutic effect of radiotherapy as a sensitizer (*3*). Recently, immune checkpoint inhibitors have been approved as first or second line therapy for unresectable, metastasized or recurrent HNSCCs (*4, 5*). However, despite these multi-modality treatments and newly developed drugs, the overall response rate remains low, with more than half of HNSCC patients dying from disease progression and treatment resistance within 2 years (*6*). Clearly, an urgent, unmet need exists to further our understanding of the pathophysiology of HNSCC to improve health care and clinical management of its patients.

We and others have demonstrated that patient-derived tumor organoids (PDOs) model the 3- dimensional (D) organization of originating tumor tissues, faithfully maintain patient-specific genomic alterations and gene expression programs, and can be *ex vivo* cultured for long term while maintaining key functional characteristics (*7–11*). Compared with the patient tumor-derived xenograft (PDX) model, the success rate of PDO culture is considerably higher. Moreover, PDOs exhibit morphological and histological similarities with the parental tumors and may reflect treatment responses of corresponding patients. Indeed, PDOs for HNSCC have been developed recently, which have demonstrated their values as important models for the development of personalized cancer therapy (*12, 13*).

While these recent important efforts in PDOs have greatly advanced the investigation of HNSCC precision medicine and predictive biomarker development, several key questions remain to be addressed: to what extent can PDOs model histopathology, drug responsiveness and mutational profiles of parental tumor specimens? Relatedly, how does long term culture affect the biology and genomic stability of PDOs? Moreover, considering the high intratumor heterogeneity of HNSCC, single-cell transcriptomic profiling is required to further delineate gene regulatory network and cellular states within the PDOs (*14, 15*). This study is designed to address these important questions by multi-omic, single-cell genomic and functional analyses of both primary tumors and matched PDO models maintained for various durations *ex vivo* (**Figure 1A**).

**Figure 1.**
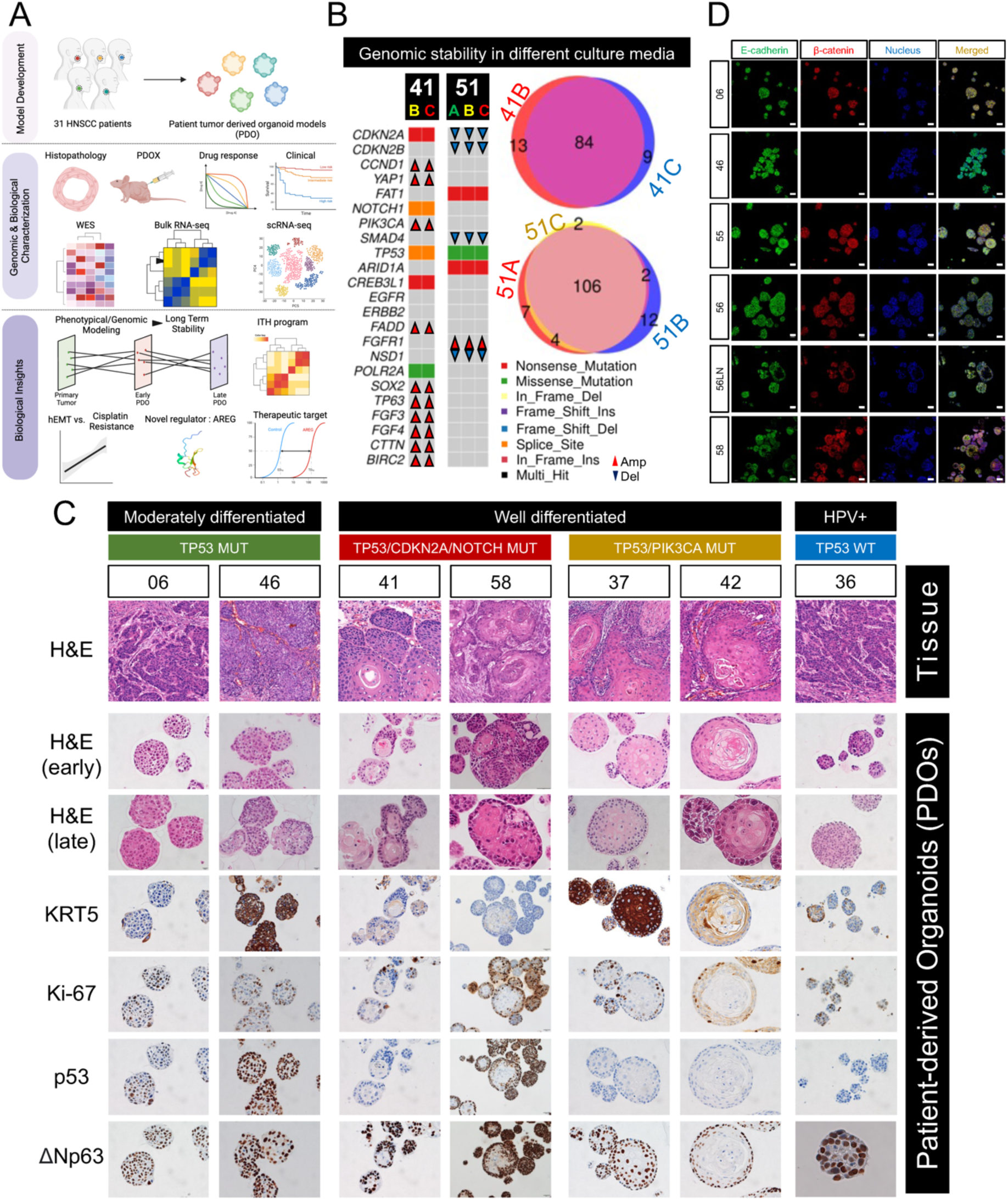
HNSCC PDOs maintain histopathological characteristics of the original tumors. **(A)** Workflow of the study design, generated by Biorender. **(B)** Oncoplots (left) of driver mutations and venn diagrams (right) of total nonsynonymous mutations from two HNSCC PDO lines grown in different culture media. (**C)** H&E and IHC staining of indicated markers on selected PDOs and their matched primary tumors. Early, early-passaged PDOs; Late, late-passaged PDOs; MUT, mutant; WT, wildtype. (**D)** The expression of epithelial markers (E-cadherin and β-catenin) were evaluated using IF staining in selected PDO lines.

## Results

### Establishment of HNSCC patient tumor-derived organoid (PDO) model

A total of 31 HNSCC patients (2 HPV^+^ and 29 HPV^-^, **Supplementary Table 1**) were enrolled in this study for the development of PDOs using surgical samples. To optimize growth conditions for tumor organoids, we prepared three different culture media based on published protocols for HNSCC, esophageal SCC (ESCC), and cervical SCC (CSCC) organoids (**Supplementary Table 2**; see Methods)(*13, 16–18*). We carefully tested every PDO line in each of the three different media independently, and grew these cultures up to 10 passages (**Supplementary Figure 1**). We then determined optimal media for each PDO line based on their growth rate (**Supplemental Table 3**). To examine the impact of different culture media on the genomic landscape, we performed whole exome sequencing (WES) on PDOs derived from two patients (**Figure 1B**). Importantly, different culture media had no significant influence on any of the genomics drivers and the mutational profile remained stable across different conditions.

After optimization, we achieved a success rate of ∼70%, and a total of 22 PDO lines were established from 21 HNSCC patients. The patient-56 had one PDO line originating from the primary tumor (PDO-56) and the other from a matched metastatic lymph node (PDO-56LN). We successfully cultured all 22 PDOs for long-term (defined as >10 passages) and characterized their morphologic features. All of the established PDOs were reproducibly recovered and expanded from cryopreservation, representing a continuous supply of viable tissues for future biological investigations. We next focused on 14 PDOs in-depth and analyzed their histopathological, genomic and intratumor heterogeneity (ITH) features, as described below.

### HNSCC PDOs retain histopathological characteristics of the originating tumors

We first performed histological analyses to compare primary HNSCC tumors *vs.* corresponding PDOs, and three major histological patterns were observed. The first pattern was well-organized, stratified squamous epithelial cells encircling an inner keratin pearl (represented by cases PDO-37 and PDO-42 in **Figure 1C**). This is reminiscent of well-differentiated squamous cell carcinoma. Indeed, both PDO-37 and PDO-42 were derived from primary tumors that were well-differentiated HNSCC, which exhibited similar histological architecture (see the Tissue panel of **Figure 1C**). The second pattern was characterized by PDOs being densely filled with basal-like, ΔNp63^+^ cancer cells (represented by cases PDO-06 and PDO-46 in **Figure 1C**), resembling moderate or poorly differentiated squamous cell carcinoma. Consistently, both 06 and 46 primary tumors were moderately differentiated HNSCC (**Figure 1C**). The last pattern had keratin pearls inside the organoid, however in contrast to the first pattern, the surrounding squamous epithelial cells were disorganized (represented by cases PDO-41 and PDO-58 in **Figure 1C**). Interestingly, this morphologic feature was only seen in HNSCC organoids having NOTCH1 mutations with concurrent mutations of TP53/CDKN2A. This is notable since a recent study on genetically engineered mouse organoid modeling of ESCC reported that only esophageal organoids introduced with TP53/CDKN2A/NOTCH1 triple mutations showed disorganized features(*19*).

Importantly, these histological features of the PDOs were maintained after long-term culture (>10 passages). As expected, Ki67 stained proliferative basal cells in the outer layer, with some of them being ΔNp63 positive (**Figure 1C**). HNSCC PDOs having missense mutations of TP53 (e.g., PDO-06, PDO-46, PDO-58) were confirmed to exhibit strong IHC staining of p53 protein. In comparison, PDOs with either TP53 frameshift mutations (e.g., PDO-37 and PDO-42), splice-site variants (e.g., PDO-41), or wild-type TP53 (e.g., PDO-36) were expectedly negative for p53 protein staining. P16, a protein biomarker for HPV^+^ HNSCC, was positively expressed uniquely in the HPV^+^ line PDO-36, which was derived from an HPV^+^ oropharyngeal tumor (**Supplementary Figure 2A**). We further performed IF staining and validated the cell surface expression of epithelial markers including E-cadherin and β-catenin proteins in HNSCC PDOs (**Figure 1D**).

Considering that organoids are propagated by adult stem cells, we additionally performed IF staining to explore the protein expression of CD44 and ALDH1A1, reported as stem cell markers for HNSCC (*20*). Indeed, the CD44 expression was observed in all HNSCC PDOs examined, while ALDH1A1 expression was more variable **(Supplementary Figure 2B**). These results demonstrate that morphological and histopathological features of primary HNSCC tumors are retained in the established PDO models, even after long term culture.

To evaluate the tumorigenicity of the HNSCC PDOs in vivo, we performed xenograft transplantation using immunodeficient mice. Two PDO lines (PDO-04 and PDO-06) were initially selected, dissociated into single cells, mixed with Matrigel, and injected into both flanks of mice (n = 3 mice for each PDO line). The transplanted HNSCC PDO xenografts (PDOX) grew successfully in the mice (4/6 and 5/6) (**Figures 2A-B**). At the end point, we collected the xenograft samples and performed H&E and IHC staining to compare their histologies with those of the parental primary tumors and PDOs. High similarity in histology and marker protein staining (including TP53 and KRT5) across the three groups was confirmed (**Figure 2C**). We also validated that these tumor cells were human origin by staining with a human nuclei antibody (**Figure 2C**). Using this protocol, we tested a total of 17 PDO lines and 8 grew successfully as PDOX, achieving a success rate of 47%.

**Figure 2.**
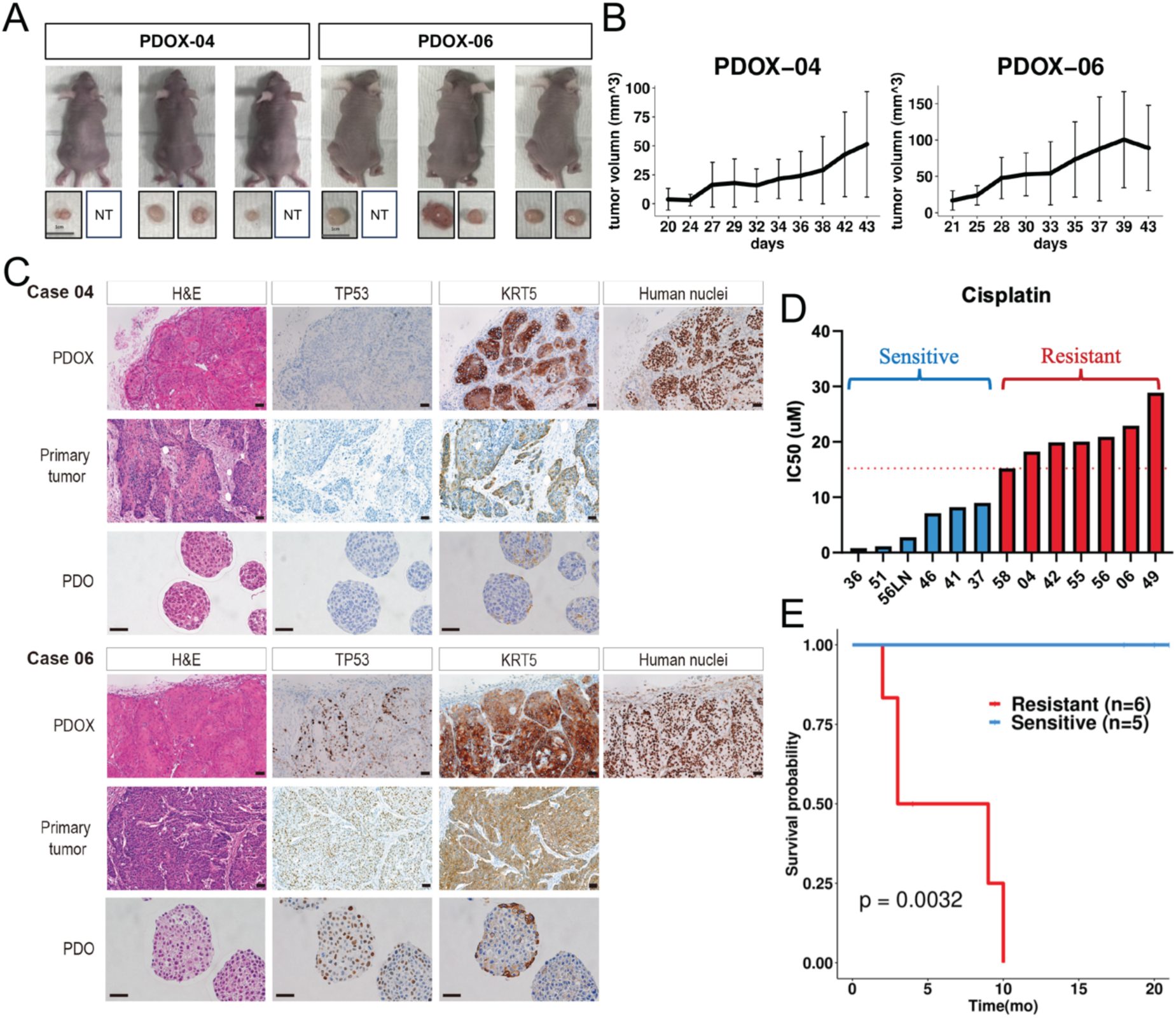
Characterization of HNSCC PDO xenograft (PDOX). **(A**) Images and (**B)** growth curves of PDOX tumors. NT, no tumor formation. (**C)** H&E and IHC staining of matched PDOX, originating tumors and PDO samples. (**D)** IC50 values of each PDO line treated with cisplatin. PDOs were divided into sensitive and resistant groups based on the median IC50 (15 uM, indicated by the dashed line). (**E)** A Kaplan meier survival plot of HNSCC patients, grouped according to the cisplatin sensitivity of their corresponding PDO lines.

### PDOs predict patient clinical response to cisplatin treatment

We next evaluated the sensitivity of PDOs to chemotherapeutics, in order to explore the potential use of organoid models as a predictive tool for treatment response. Cisplatin, a chemotherapeutic drug most commonly used in either definitive or adjuvant settings for HNSCC patients, was tested. In vitro assays showed variable responsiveness of PDOs to cisplatin (**Figure 2D**). We determined half-maximal inhibitory concentration (IC50) of each PDO line, and stratified them into resistant (> median IC50) or sensitive (< median IC50) groups (**Figure 2D**). The drug responsiveness of PDOs was then correlated with clinical outcomes of matched HNSCC patients who underwent cisplatin-based treatment strategies (surgery followed by adjuvant chemoradiotherapy using cisplatin) having a sufficient follow-up of more than 18 months. Importantly, we observed a striking association between the drug sensitivity of PDOs with clinical outcome of matched patients. Specifically, the disease-free survival of patients whose PDOs were resistant to cisplatin was significantly lower than that of patients whose PDOs were sensitive (p = 0.0032, **Figure 2E**). This result strongly suggests that HNSCC PDOs are capable of predicting patients’ clinical response to cisplatin-based standard treatment.

### PDOs preserve the genomic landscape of parental tumors

To determine the extent to which the PDOs maintain the mutational landscape of the parental tumors, WES was performed on the “trio” set, including early- and late-passaged PDOs as well as matched tumor samples from 13 patients, using paired blood DNA as the germline control. The overall mutational pattern of PDOs was highly consistent with that of primary tumors (**Figure 3 and Supplementary Figure 3**). All of the PDOs except for PDO-57 contained somatic mutations in two or more established cancer driver genes, such as those frequently mutated in HNSCC: TP53 (62%), CDKN2A (27%), FAT1 (31%), PIK3CA (15%), NOTCH1 (15%), KMT2D (8%), CAPS8 (8%), KEAP1 (8%) (**Figures 3A-B**), confirming the malignant nature of these PDOs. Since PDO- 57 did not contain any matched driver mutations, it was not included in the following analyses (**Supplementary Figure 4**). We mapped these gene mutations to genomically-altered signaling pathways curated by The Cancer Genome Atlas (TCGA), confirming that mutations in the TP53 pathway were the most frequent (62%), followed by the Hippo pathway (54%), cell cycle pathway (25%) and NOTCH pathway (25%) (*21*). Other recurrantly mutated pathways included the WNT pathway (21%), PIK3CA pathway (17%), and NRF2 pathway (8%). Importantly, the mutational landscape showed a high concordance between primary tumors and matched early-passaged PDOs (average overlapping rate=76%, **Figure 3C**). Moreover, the mutational profile of early-passaged PDOs was also stably preserved in the late-passaged cultures (average overlapping rate=87%, **Figure 3D**).

**Figure 3.**
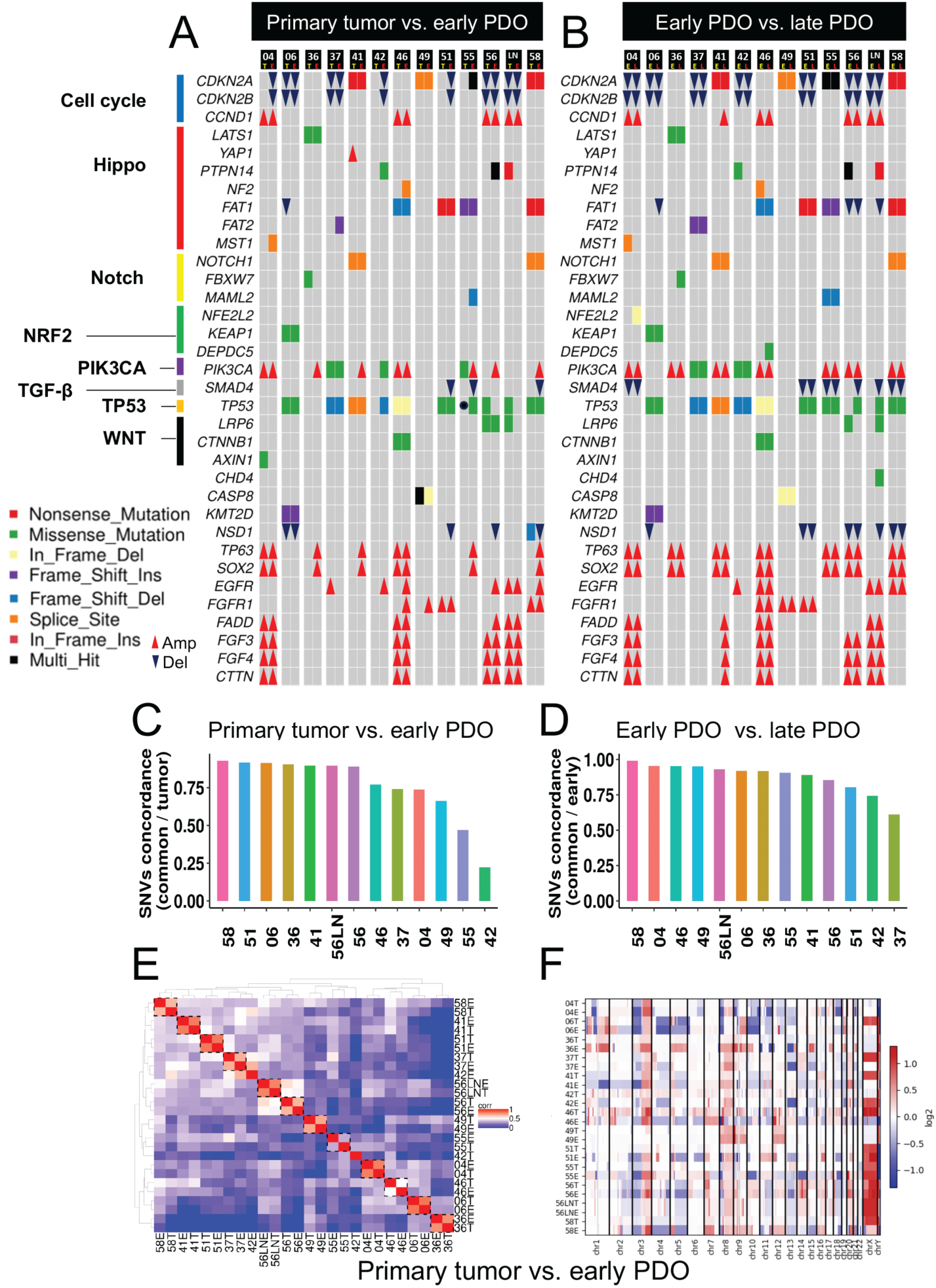
HNSCC PDOs preserve the genomic landscape of the parental tumors. **(A)** Oncoplots of driver mutations and copy number variations (CNVs) comparing parental tumors (T) *vs.* early-passaged PDOs (E), or **(B)** between early- and late-passaged PDOs (L). TCGA-curated genomic pathways are annotated on the left. **(C)** Column plots summarizing the concordance rate of total nonsynonymous mutations between parental tumors *vs.* early-passaged PDOs, or **(D)** between early- and late-passaged PDOs. (**E)** Heatmap showing the unsupervised clustering of parental tumors and early-passaged PDOs. (**F)** CNV profiles of parental tumors and early-passaged PDOs.

We next calculated mutational variant allele frequency (VAF) in the original tumors and PDOs. The VAF values of early-passaged PDOs were significantly and strongly correlated with that of parental tumors (average Pearson correlation R=0.79, **Supplementary Figure 5A and Supplementary Table 4**). Not surprisingly, the VAF values of PDOs were always higher than that of original tumors, due to non-malignant cell contamination in the primary samples. In addition, the VAF values of early- and late-passaged PDOs correlated strongly and consistently (average Pearson correlation R=0.93, **Supplementary Figure 5B and Supplementary Table 4**).

Despite the overall high genomic consistency between PDOs and their corresponding tumors, we did notice a discordant mutational profile in one patient, T42. Specifically, somatic mutations in driver genes (e.g., *TP53, PIK3CA, DCC*) detected in PDO-42 were not observed in the matched primary tumor sample T-42. Since these driver mutations were consistently identified in both early- and late-passaged PDO-42 lines, we speculated that their absence in T-42 was due to the low tumor purity of the primary tumor, which could hinder the detection of mutations with low VAF. We thus measured the malignant cellularity of each sample by the Sequenza method (*22*). Indeed, T-42 had the lowest tumor purity among all primary tumors. As expected, all of the PDO lines showed near 100% tumor purity, except for PDO-42 (69%), which was still much higher than most of the primary samples (**Supplementary Figure 6A**).

We next analyzed copy number alterations (CNA) using the CNVkit package (*23*). As anticipated, we detected recurrent copy number losses of 3p and 8p, and copy number gains of 3q, 5p, and 8q chromosome regions **(Figure 3F**), which are known to be prevalent in HNSCC primary tumors (*24*). Many HNSCC-relevant focal CNAs were also confirmed, such as deletions of *CDKN2A, NSD1, SMAD4*, and amplifications of *TP63, SOX2, PIK3CA, CCND1, EGFR and MYC* (**Figures 3A-B**). Importantly, Pearson correlation analysis followed by unsupervised clustering demonstrated highly similar CNA profiles between parental tumors and matched early-passaged PDO lines (**Figure 3E**). Moreover, this strong similarity was also retained between early- and late-passaged PDO lines **(Supplementary Figure 6B**). These results suggest that PDOs faithfully maintain the genomic landscape of the originating tumors, including both somatic mutations and CNAs.

### Bulk transcriptomic analysis of PDOs

To understand gene expression patterns of these tumor organoids, we performed bulk RNA-seq on all 13 early-passaged PDOs which had matched WES data. Using geneset signatures established for HNSCC molecular subtypes (namely classical, basal and atypical) (*25*), we calculated the GSVA scores to determine the molecular subtype of each PDO. The majority PDO lines had a dominating subtype assignment (**Figure 4A**), which were separated by the PCA visualization with certain overlap (**Figure 4B**). PDO-36, an HPV+ tumor line, was expectedly classified as atypical. PDO-56LN was derived from a metastatic lymph node, and exhibited a different subtype (atypical) than PDO-56 (basal) which was from the paired primary tumor (**Figure 4A**), suggesting intra-patient tumor heterogeneity.

**Figure 4.**
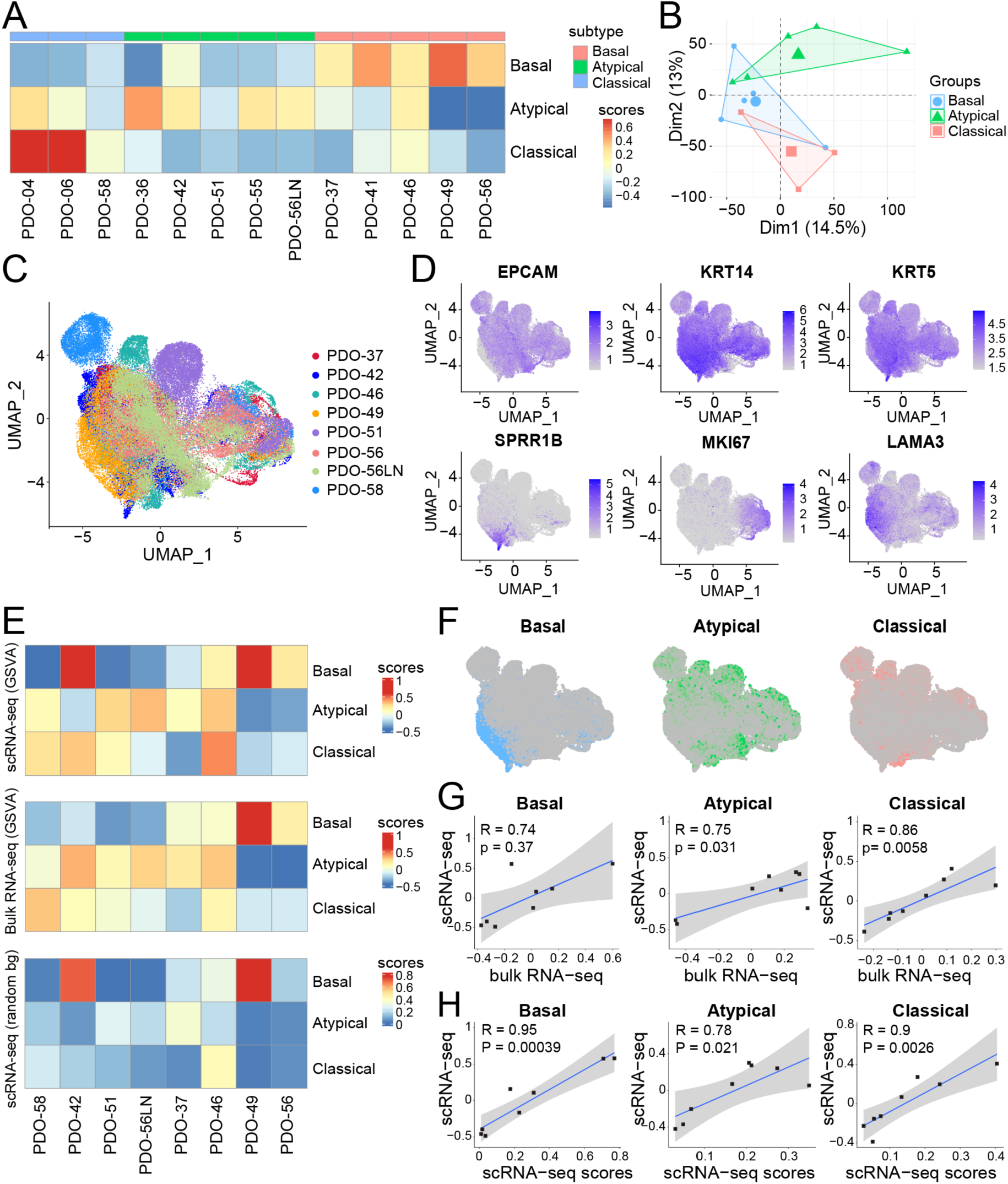
Bulk and single-cell transcriptomic analysis of HNSCC PDOs. **(A)** Heatmap and (**B)** PCA plot showing molecular subtyping of 13 HNSCC PDOs based on bulk RNA-seq. (**C)** A UMAP showing 68,919 cells obtained from 8 PDO lines, colored by patient origin. (**D)** UMAP of the expression of selected marker genes. (**E)** Heatmap showing consistent molecular subtyping of PDOs using either scRNA-seq or bulk RNA-seq. (**F)** UMAP of subtype annotation of single cells. (**G)** Pearson correlation of subtype scores derived by scRNA-seq *vs*. bulk RNA-seq. (**H)** Pearson correlation of subtype scores derived by two independent methods based on scRNA-seq.

### Single-cell RNA-seq profiling of PDOs

Intratumoral heterogeneity (ITH) and functional diversity of tumor cells is a cancer hallmark, enabling tumor immune evasion, metabolic adaptation, and resistance to treatment. Since our above data and others (*13, 16–18*) have demonstrated that PDOs preserve patient-specific genomic and transcriptomic profiles and maintain pathological characteristics of corresponding tumors, we performed single-cell RNA-seq (scRNA-seq) to delineate cellular plasticity and intratumoral expression heterogeneity of 8 HPV^-^ PDOs, which also had matched RNA-Seq and WES data. After sequencing quality control and standardization, a total of 68,919 cells were obtained using a droplet-based system enabling 3′ mRNA counting (average=8,615 cells/PDO). An average of 23,742 (22,202 ∼24,606) genes were measured in each cell (**Figure 4C**).

Using established cell-type specific markers, we first confirmed that our PDOs almost exclusively comprised squamous epithelial cells (e.g., KRT14, KRT5, EPCAM, **Figure 4D**) and few stromal cells were detected (**Supplementary Figure 7A**). UMAP plotting showed that these organoid tumor cells were clustered primarily by patient identity, liekly due to patient-specific genomic drivers, consistent with previous findings (*14, 26*). Considering that molecular subtypes likely also contribute to the transcriptional diversity among PDO lines, we determined molecular subtypes of each single cell using the GSVA method. Notably, scRNA-seq derived molecular subtypes were highly consistent with those determined by bulk RNA-seq from matched samples (**Figure 4E**) in all cases but PDO-42, which is possibly due to moderate tumor cell content (69%), as we described before (**Supplementary Figure 6A**). Indeed, this overall high correlation was confirmed by Pearson correlation analysis (**Figure 4G**). To validate further this result, we calculated the molecular subtype enrichment score for every single cell using an independent method, based on the normalization and ranking using random genesets (See Method). Importantly, this random geneset approach was highly consistent with the GSVA results and again strongly correlated with bulk RNA-seq data (**Figures 4E, H**).

### Intratumor transcriptional heterogeneity (ITH) of PDOs

Recent work has identified gene modules (i.e., sets of coordinately expressed genes) as the defining features of cellular states (*14, 27–29*). To decipher individual cancer-cell states and transcriptional ITH of PDOs, we performed non-negative matrix factorization (NMF) to identify gene modules as described previously (see Methods) (*15, 28–30*). This integrative computational approach attempts to alleviate the impact of both technical confounders (e.g., batch effects) and patient-specific characteristics, which often hinder the goal of identifying common expression programs across different patient tumors. As a result, we identified eight clusters of recurrent gene modules across individual PDOs (**Figure 5A**). For each of the clusters, based on their shared genes, we defined 8 consensus gene programs, each with genes coordinately upregulated by subpopulations of cells in at least 2 PDO lines, hence representing heterogeneously expressed intratumoral programs (henceforth “ITH programs”).

**Figure 5.**
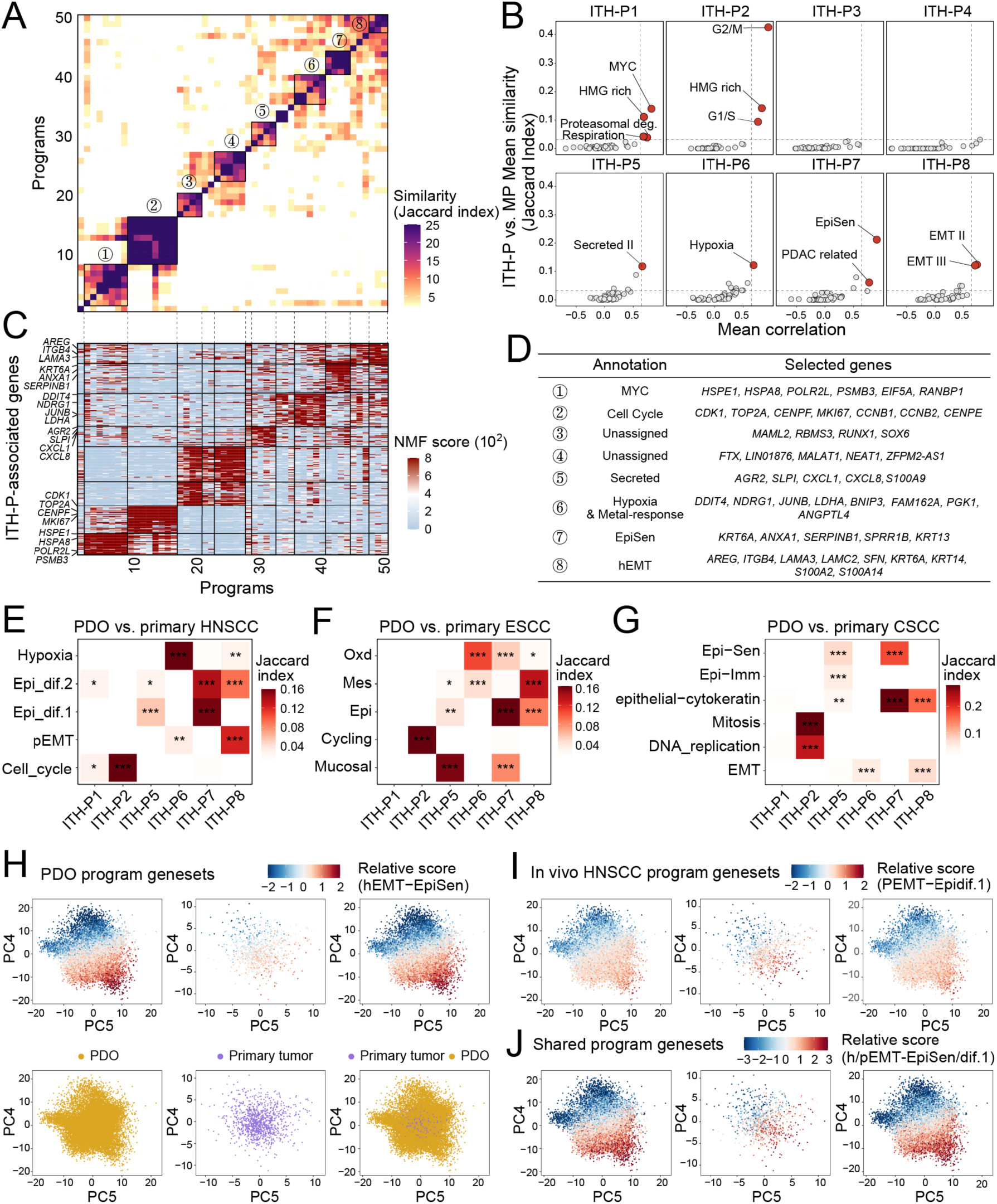
scRNA-seq reveals the intratumor heterogeneity (ITH) of HNSCC PDOs. **(A)** Heatmap showing unsupervised clustering of 50 NMF programs based on top program genes, with similarity measured by Jaccard indices. Eight clusters of consensus ITH programs are numbered and highlighted. (**B)** Dot plots showing the Jaccard index (y) and mean correlation (x) between ITH programs and published meta-programs (*15*). Dashed lines indicate a 99.9% confidence threshold determined using permutations of NMF programs. (**C**) A heatmap showing expression levels of program genes, with each program aligned to panel (**A)**. (**D)** The most significantly correlated ITH programs from panel (**B)**. (**E-G)** Heatmaps showing similarity (measured by Jaccard indices) between PDO ITH programs and published programs from primary HNSCC (**E**), ESCC (**F**) and CSCC (**G**). *, FDR q < 0.05; **, FDR q < 0.01; ***, FDR q < 0.001; hypergeometric test. (**H)** PCA plots of PDO cells only (yellow, lower left), primary HNSCC tumor cells only (purple, lower middle) or combined (lower right). In the upper panel, cells are colored by the relative score for the hEMT minus EpiSen program identified from PDO samples. **(I-J)** similar PCA plots to panel H, except using program genesets identified from HNSCC primary tumors **(I)** or shared program genesets between PDO and primary tumors **(J)**.

To understand these eight ITH programs, we annotated them by performing enrichment analysis against well-curated programs established by the same NMF approach from large-scale scRNA-seq delineation of pan-cancer tumor samples (*15*). The ITH programs in our PDO models were annotated to MYC (Program-1), Cell cycle transition (Program-2), Secreted (Program-5), Hypoxia (Program-6), Epithelial senescence (EpiSen, Program-7), shown in **Figures 5B-C**. For the last Program-8, while pan-cancer epithelial-mesenchymal transition (EMT) II/III program genes were significantly enriched, this program contained many keratinization-related genes, such as SFN, KRT6A, KRT14, S100A2, S100A14 (**Figure 5D, Supplementary Table 5**), suggesting a hybrid epithelial/mesenchymal state. Indeed, program-8 resembled both EMT/mesenchymal programs and epithelial differentiation programs across different types of SCC (**Figures 5E-G**), and thus we annotated Program-8 as a hybrid EMT (hEMT) program.

Importantly, these ITH programs resembled strongly and significantly (based on Jaccard index and p value) scRNA-seq-derived *in vivo* intratumoral programs from published HNSCC patients (**Figure 5E**) and other SCC patients such as ESCC (**Figure 5F**) and CSCC (**Figure 5G**), suggesting that our PDOs faithfully model intratumor expression heterogeneity pattern of *in vivo* patient tumor samples (*14, 29–31*).

To further ascertain that ITH programs of our PDO models recapitulate those expressed in HNSCC patient samples, we performed combined analysis of our organoid data together with *in vivo* scRNA-seq data of HNSCC fresh tumors (*14*). Indeed, PCA plots showed that our PDO cells and *in vivo* HNSCC primary tumor cells were intermingled and occupied similar expression space (**Figure 5H**, bottom panel), suggesting an overall transcriptional comparability between these two models. We next plotted relative scores between the EpiSen (ITH Program-7) and the hEMT program (ITH Program-8) on each single cell. We chose these two programs because prior studies have demonstrated that EMT and EpiSen represent two polarized cellular states in subsets of intratumoral malignant cells (*32*). As anticipated, in our PDO samples, hEMT^+^ and EpiSen^+^ cells were completely separated and resided in two extreme regions (**Figure 5H**, top left). Importantly, these ITH program genes established by our scRNA-seq approaches also effectively identified and separated hEMT^+^ and EpiSen^+^ cells in *in vivo* HNSCC primary tumors (**Figure 5H**, top middle).

We next performed a reciprocal analysis, i.e., using *in vivo* program genes established by scRNA-seq of patient tumors, we plotted the relative scores on our PDO cells. Importantly, we confirmed that *in vivo* program genes also clearly separated hEMT^+^ and EpiSen^+^ states in our PDO cells (**Figure 5I**). Moreover, shared program genes between our *ex vivo* PDO samples and *in vivo* HNSCC primary tumors also effectively separated these two cellular states (**Figure 5J**). These results together suggest that ITH programs identified by PDO scRNA-seq faithfully recapitulate cellular states and intratumoral transcriptional heterogeneity of *in vivo* patient tumors.

### An hEMT ITH program is strongly associated with cisplatin resistance of PDOs

Interestingly, while cell cycle programs were expectedly observed in every PDO line, the fraction of other ITH program cells, such as hEMT, varied markedly between PDOs, from as little as 4.36% in PDO-51 to as much as 97.4% in PDO-49 (**Figures 6A-B, Supplementary Figure 7B**). We next asked whether any of the ITH programs might contribute to the variation of drug responsiveness we observed earlier (**Figure 2D**). Our 8 scRNA-seq profiled PDOs had 4 cisplatin-sensitive and 4 cisplatin-resistant lines based on their median IC50 value defined earlier, and we compared the summarized score of each ITH program between these two groups by a student t test. Interestingly, across all ITH programs, only the hEMT program showed statistically significant differences between the sensitive and resistant groups. Specifically, the hEMT program had a higher score in the resistant than sensitive group (**Figure 6C**). Consistently, the subpopulation ratio of hEMT^+^ cells was also significantly larger in the resistant than sensitive PDOs (**Figure 6D**). Strikingly, Pearson correlation analysis showed that both the hEMT program score (R=0.9, P=0.0022) and hEMT^+^ cell ratio (R=0.91, P=0.0019) were prominently and significantly correlated with the IC50 value of PDOs (**Figures 6E-F**).

**Figure 6.**
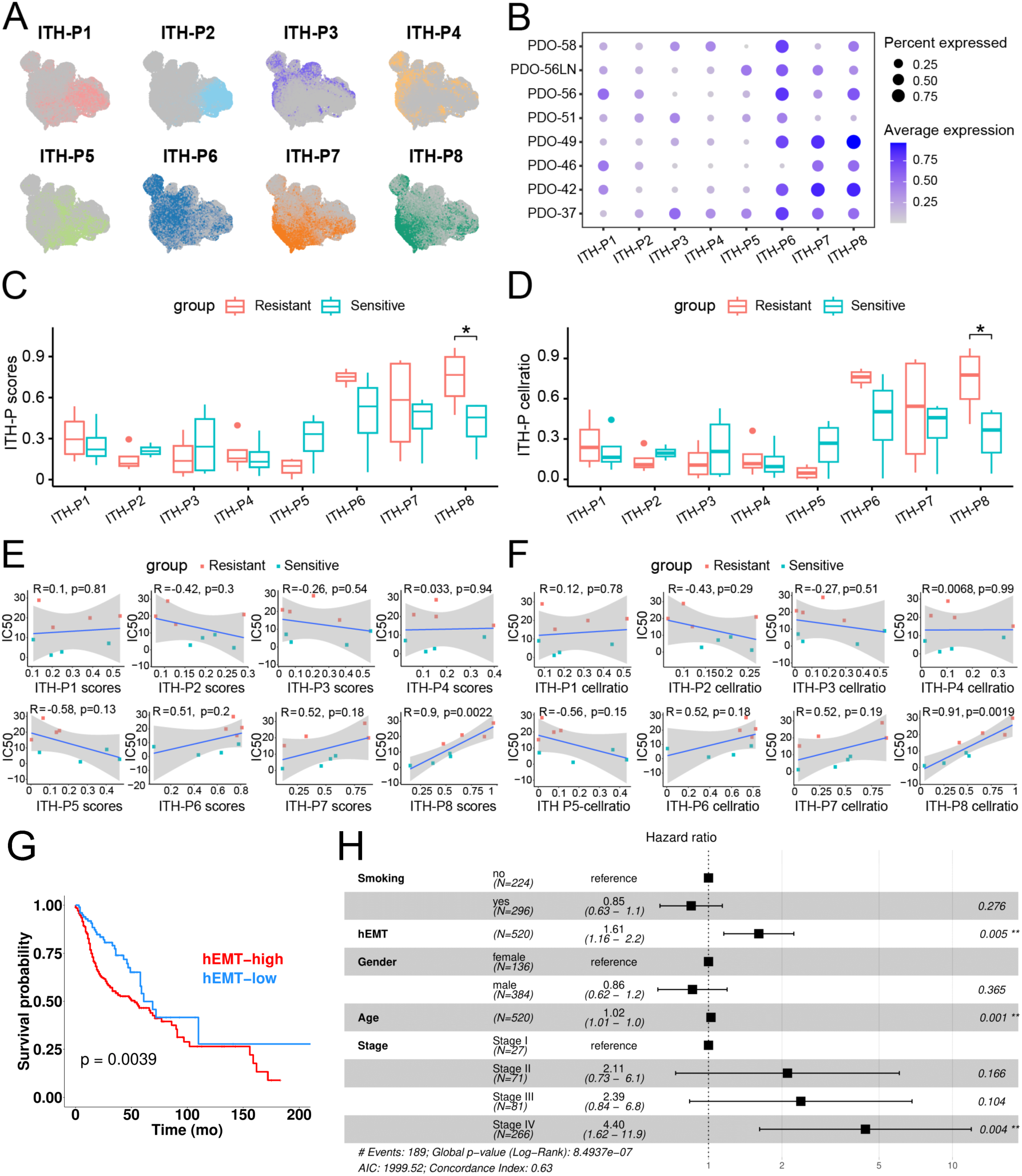
The hEMT program is associated with PDO cisplatin resistance. **(A)** UMAPs showing the distribution of each ITH program at the single-cell level. (**B)** Dot plots showing the program score and ratio of each ITH program in each PDO sample. (**C-D)** Box plots showing the program score (**C**) and cell ratio (**D**) of each ITH program comparing the cisplatin-sensitive group (n=4) with the resistant group (n=4). (**E-F)** Pearson correlation between program score (**E**) and cell ratio (**F**) of each ITH program *vs.* IC50 values of cisplatin. Each dot is one PDO sample. (**G)** A Kaplan meier survival plot of TCGA HNSCC patients (n=520), grouped according to the mRNA expression of 32 hEMT genes using the GSVA method. (**H)** multivariate survival analysis of indicated factors using the TCGA HNSCC cohort.

Resistance to cisplatin treatment is a well-recognized prognostic factor for HNSCC patients (*27, 28*), which was also recapitulated by our PDO models, as described earlier in **Figure 2E**. We thus asked whether the hEMT program was associated with HNSCC patient survival by analyzing the TCGA HNSCC cohort. Specifically, we scored the hEMT program gene signature (n=32 genes, **Supplementary Table 6**) by the GSVA method using the TCGA bulk RNA-seq data. Importantly, patients with hEMT^-high^ tumors had significantly worse overall survival compared with those with hEMT^-low^ tumors (**Figure 6G**). Moreover, a multivariate analysis confirmed that the hEMT program score is an independent prognostic factor for HNSCC patients (**Figure 6H**).

We next inspected the 32 hEMT program genes in depth, and first confirmed that the majority of them were upregulated in the resistant vs. sensitive group (**Supplementary Figure 7C**). Moreover, the expression of most of these genes showed positive correlation with the IC50 values of PDO lines (**Figure 7A**).

**Figure 7.**
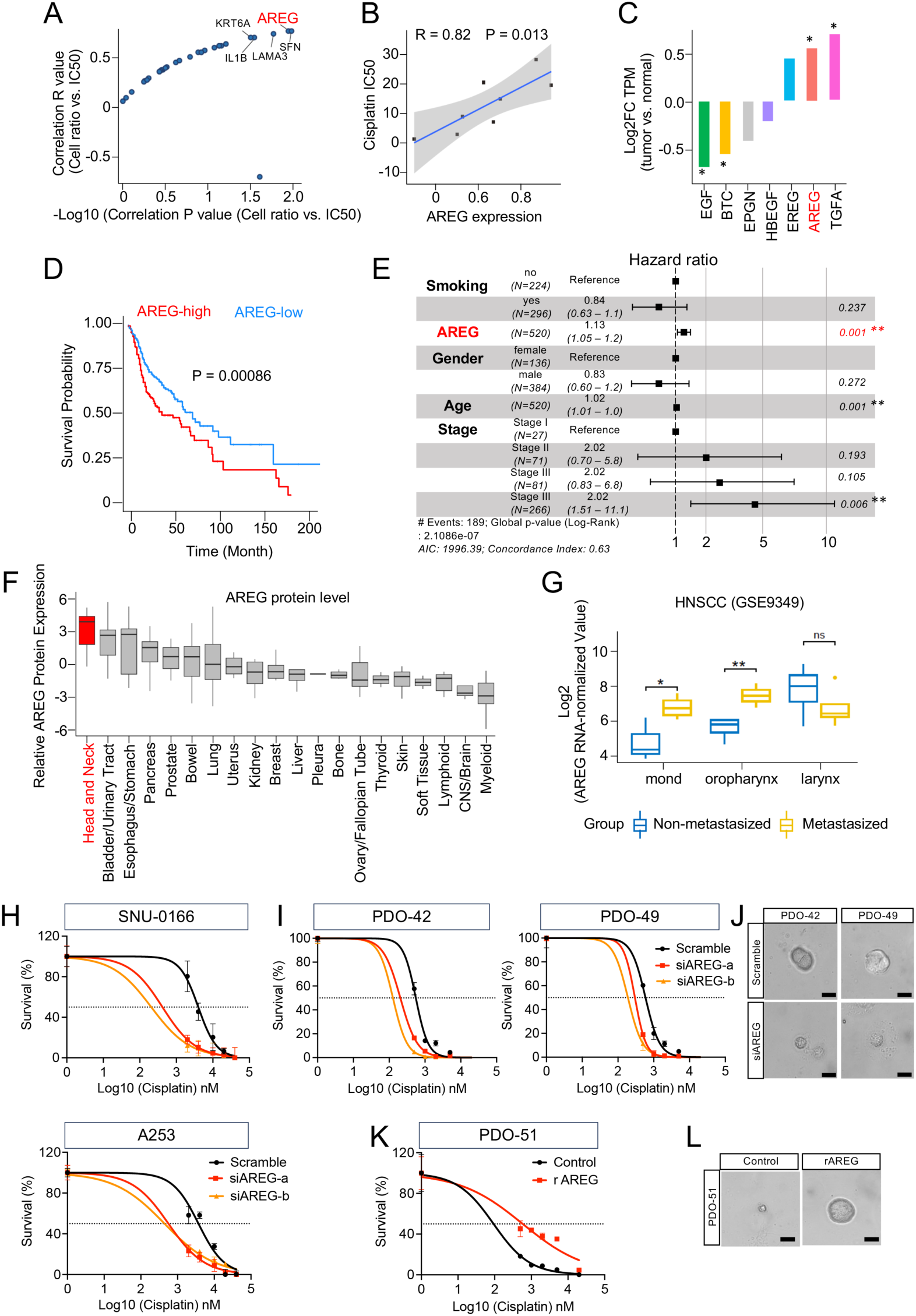
AREG, a top hEMT program gene, regulates PDO responsiveness to cisplatin. **(A)** A scatter plot showing the Pearson correlation R value (y) and p value (x) of each hEMT gene with IC50 values of cisplatin. Top 5 genes are highlighted. (**B)** Pearson correlation between AREG expression and IC50 values of cisplatin. Each dot is one PDO sample. (**C)** A column plot showing the expression fold change of seven EGFR ligands between HNSCC tumors with normal samples using TCGA bulk RNA-seq data. *, P < 0.05. (**D)** A Kaplan meier survival plot of TCGA HNSCC patients (n=520), grouped according to the mRNA expression of AREG. (**E)** multivariate survival analysis of indicated factors using the TCGA HNSCC cohort. (**F)** A box plot showing the protein levels of AREG across various HPA human tumor samples. (**G**) Box plots showing the mRNA expression of AREG in indicated HNSCC tumor samples from the dataset GSE9349. (**H)** dose-response curves of scramble control and AREG-knockdown samples to cisplatin treatment in HNSCC cell lines (A253 and SNU-1066) or (**I)** PDOs (PDO-42 and PDO-49). (**J)** Images of PDOs upon cisplatin treatment. (**K)** dose-response curves and (**L)** images of PDO-51 to cisplatin treatment in the presence and absence of recombinant AREG protein. Scale bar; 40 μm

### AREG regulates hEMT ITH program and cellular sensitivity to cisplatin treatment

We were particularly interested in the AREG gene, ranked the 2nd highest in terms of its expression correlation with cisplatin IC50 (**Figures 7A-B**), because it was not only upregulated in HNSCC tumors vs adjacent nonmalignant samples (**Figures 7C**) but also was significantly associated with HNSCC patient survival in Kaplan-Meier survival analysis (**Figure 7D**) and multivariate cox regression analysis (**Figure 7E**). The AREG gene encodes a growth factor (amphiregulin) which acts as a low-affinity ligand for EGFR, a key druggable driver for HNSCC tumors (*33, 34*). Interestingly, among all seven EGFR ligands, only AREG and TGFA were significantly overexpressed in HNSCC tumors (**Figure 7C**). Notably, among all human cancers, HNSCC had the highest AREG protein level based on pan-cancer proteomic analyses by the HPA project (**Figure 7F**) (*35*). In addition, among all seven EGFR ligands, AREG showed the strongest correlations with the EGFR pathway activity, as inferred by the PROGENy method, reported by a recent large-scale HNSCC proteomics study (*33*). Moreover, AREG was expressed higher in metastasized than non-metastasized primary HNSCC (**Figure 7G**).

Prompted by these intriguing findings, we decided to experimentally test if AREG functionally contributes to cisplatin resistance and EGFR pathway activation in the context of HNSCC cancer biology. We first selected HNSCC cell lines with high AREG baseline expression level (**Supplementary Table 6**) and performed loss-of-function assays. Importantly, knockdown of AREG by independent siRNAs or shRNAs enhanced cellular sensitivity to cisplatin treatment (**Figure 7H, Supplementary Figure 8**). More importantly, we repeated this loss-of-function assay in our resistant PDO lines, and observed that AREG knockdown re-sensitized these PDOs to cisplatin (**Figures 7I-J**). Conversely, incubation of recombinant AREG protein conferred resistance of our HNSCC PDOs to cisplatin treatment (**Figures 7K-L**).

We next performed protein assays and observed that silencing of AREG reduced the phosphorylation of both AKT and ERK, central downstream targets of EGFR signaling in HNSCC (**Figure 8A**). Interestingly, we found very strong expression correlation between AREG and many other hEMT program genes in both TCGA and HPA datasets (**Figures 8B-C**), implying that AREG may functionally contribute to the regulation of the hEMT program activity (*24, 35, 36*).

**Figure 8.**
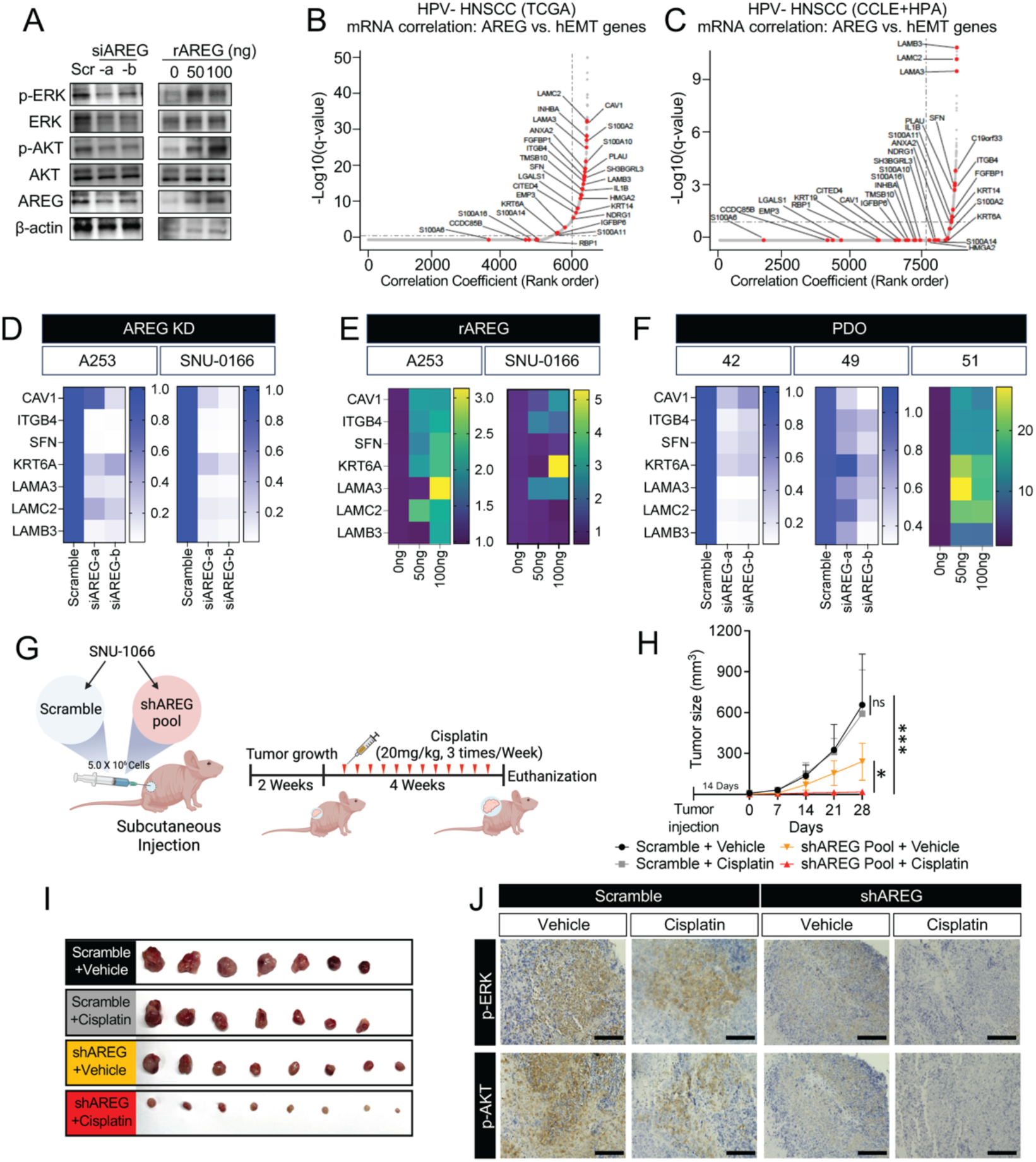
AREG regulates the activity of hEMT program in HNSCC. **(A)** Western blotting of indicated antibodies upon either knockdown of AREG with siRNAs or addition of recombinant AREG protein in SNU-1066 cells. (**B-C)** hockey stick plots showing the correlation of the mRNA expression of AREG with other genes, rank ordered by correlation coefficient. hEMT program genes are highlighted in red. Only genes with nonnegative correlation are shown. (**B)** is the TCGA dataset and (**C)** is CCLE+HPA combined dataset. (**D-F)** heatmaps showing changes in expression of hEMT program genes upon either AREG knockdown, or addition of recombinant AREG protein, in HNSCC cell lines or PDO samples. (**G**) A schematic graph of the experiment design, created using Biorender. (**H**) Growth curves and (**I**) images of xenograft tumors. **(J)** Representative IHC images of p-ERK and p-AKT on each group. Scale Bar; 100 μm

We thus measured the expression of hEMT program genes exhibiting high correlations with AREG in the absence or presence of AREG knockdown. Importantly, depletion of AREG down-regulated the levels of hEMT program genes (**Figure 8D**). Reciprocally, addition of recombinant AREG protein promoted the expression levels of these hEMT genes (**Figure 8E**). These results were also reproduced in HNSCC PDO models (**Figure 8F**). Finally, we extended these findings to an in vivo setting by performing a xenograft assay evaluating the function of AREG in cisplatin resistance (**Figure 8G**). Specifically, we established scramble control or AREG-silenced SNU-1066 xenograft tumors followed by the treatment of cisplatin. While AREG knockdown itself reduced tumor growth, cisplatin treatment completely blocked tumor development in AREG-silenced group. In contrast, control tumors were almost unresponsive to cisplatin treatment (**Figures 8H-J**). Taken together, these results demonstrate that AREG, a top hEMT program factor, functionally contributes to the cellular resistance of cisplatin treatment, possibly due to the activation of EGFR pathway.

## Discussion

Developing a preclinical model capable of accurately maintaining patient-specific tumor characteristics is imperative for the advance of precision cancer medicine. In this study, we present a successful culture of HNSCC tumor organoids using patient surgical specimens. Our sustained maintenance of these PDOs for extended durations validates the preservation of crucial histopathological and genomic features from the original patient tumors within the organoids.

Previous research has noted varying success rates, typically between 30% to 80%, in generating HNSCC tumor organoids, which modestly fall behind the success rates observed in gastrointestinal (GI) malignancies (*37–40*). While precise reasons remain incompletely understood, factors such as cell-type specificity and inter-patient heterogeneity are likely influential in determining organoid development efficiency. By carefully optimizing organoid culture media (*13, 16–18*), we have established PDOs from both oral and pharyngeal tumors, encompassing both HPV+ and HPV-cancer tissues, achieving an overall success rate of approximately 70%.

While prior studies generally suggest that patient-specific genomic abnormalities can be preserved in PDOs, a trio comparison between parental tumors, early- and late-passaged organoids has rarely been performed. Our WES analyses have confirmed that despite long-term culture, the mutational landscape, VAF values and CVA profiles of primary tumors are robustly preserved in early-passaged PDOs, and stably maintained in late-passaged cultures. Likewise, histopathological characteristics of parental tumors were also retained in both early- and late-passaged PDOs. We successfully created a xenograft model using our HNSCC organoids and confirmed that the morphology of the formed tumor showed high similarity to both the parental tumor and PDOs.

The presence of diverse cellular states within each tumor underlies intratumor heterogeneity (ITH), posing a significant challenge in cancer therapeutics. A number of recent studies have embarked on delineating ITH through scRNA-seq of fresh tumor specimens (*41*). An emerging concept gleaned from prior scRNA-seq studies is the identification of expression modules associated with ITH, which encompass sets of genes exhibiting coordinated expression variability across single malignant cells within a specific tumor. To our knowledge, the present work is the first to systematically delineate ITH programs using scRNA-seq of tumor organoids. Importantly, integrative analyses confirmed that ITH programs identified from our PDOs resembled strongly and significantly those identified from *in vivo* HNSCC patient samples, suggesting that PDOs robustly model diverse cellular states and intratumoral heterogeneity of primary tumors. This finding holds important implications since the majority of ITH studies rely on static, descriptive tumor sequencing. The capability of PDO in modeling ITH programs enables functional interrogation of diverse intratumoral cellular states within a given tumor.

Indeed, here we discovered a prominent correlation between the hEMT ITH program and cisplatin resistance in HNSCC PDOs. We further showed that the hEMT program score is an independent prognostic factor for HNSCC patients. A similar partial EMT (pEMT) program was discovered by scRNA-seq profiling of HNSCC tumors (*14*). Importantly, our hEMT program is distinct from the classical EMT phenotype since most EMT-related transcription factors are not expressed in this unique cellular state. Moreover, the hEMT ITH program is also distinguished from the canonical EMT since it expresses both epithelial and mesenchymal genes, indicating an epithelial/mesenchymal hybrid state. Indeed, our hEMT program contains a number of genes associated with keratin differentiation, including SFN, KRT6A, KRT14, S100A2, S100A14, in addition to common EMT factors such as INHBA, ITGB4, LAMA3, LAMC2 (**Supplementary Table 5**).

Our study unveils amphiregulin (AREG), a top factor within the hEMT ITH program, as a key contributor to cisplatin resistance in HNSCC. Functional experiments involving AREG loss/gain-of-function in cell lines, PDOs and clinical samples consistently demonstrated a role of AREG in mediating chemoresistance via EGFR pathway activation. This finding aligns with previous observations identifying AREG’s involvement in drug resistance (*42–45*). However, while our study predominantly focused on AREG, it is crucial to recognize that the hEMT ITH program may encompass various genes regulating cellular plasticity and treatment responses. Our singular exploration of AREG raises the possibility that other genes regulating the hybrid epithelial/mesenchymal states might also contribute to cisplatin resistance. This observation prompts the need for further comprehensive investigations into the collective impact of multiple hEMT genes on drug responsiveness in HNSCC.

## Materials and Methods

### Sample collection and tissue processing

Written informed consent for sample collection and research use was obtained from all 31 patients. The Institutional Review Board of Yonsei University approved the conduct of this study (IRB number: 3-2022-0128). Fresh tumor tissues and matched peripheral blood were collected from HNSCC patients who underwent surgery at the Severance Hospital. The clinico-pathological data of patients who participated in this study are summarized in **Supplementary Table 1.**

Tissue samples were collected from surgical specimens and divided into multiple pieces for DNA/RNA extraction, paraffin block preparation and organoid culture. For the organoid culture, 25 mg of the tumor tissue was finely minced and placed in DMEM/F12 medium containing 1 ml of 5 mg/ml collagenase II and 10 μM Y-27632 (ROCK I/II inhibitor). Then, it was kept in an incubator at 37° C for 1 hour. DMEM/F12 medium was added to neutralize the enzyme, and then the samples were centrifuged, and the supernatant was removed. Subsequently, 1 ml of TrypLE with 10 μM Y-27632 was added, and the samples were resting in an incubator at 37°C for 10 minutes. After filtering residual tissue debris using a 70 μm strainer, 2 x 10^4^ cells were prepared and resuspended in 40 μl of Matrigel in each well of a 24-well plate. Next, the organoid culture medium was filled into each well. When the organoids had formed and growth was visible, subcultures were performed at intervals of 10-14 days.

### Cell lines and culture

Human HNSCC cell line, A253 was obtained from the ATCC, and SNU-1066 was purchased from Korean Cell Line Bank. A253 and SNU-1066 cell lines were cultured in McCoy’s 5A medium (Corning, Cat#10-050-CV) and RPMI medium (Corning, Cat#10-040-CM), respectively. All media were supplemented with 10% Fetal Bovine serum (FBS), (Omega Scientific, Tarzana, USA, Cat#FB-02) and 1% penicillin-streptomycin sulfate (Thermo Fisher Scientific, #30-002-CI). Cell cultures were maintained in a 37 °C incubator with 5 % CO2. Both cell lines were tested for mycoplasma contamination. To knockdown of AREG, we transfected two different siRNAs (IDT technologies) or shRNA viral particles (Santa Cruz Biotechnology, Cat#sc-39412-SH) using transfection reagent BioT (Bioland Scientific, Cat#B01-01). To increase AREG protein level in the cell lines, we added 50 ng and 100 ng of recombinant AREG protein (R&D Biotechne, Cat#262-AR-100/CF) and incubated for 72 hours. To investigate the chemoresistance of cisplatin upon the AREG expression levels, we treated multiple concentrations ( 0, 0.5, 1, 2, 5, and 20μM) of cisplatin (Selleck Chemical, Cat#NC0564900) to the cell lines for 72 hours.

### Tumor organoid culture

Tumor organoid culture medium included the following niche factors: B27, Noggin (TGF-β inhibitor), Y27632 (ROCK I/II inhibitor), nicotinamide, SB202190, A83-01 (TGF-β inhibitor), forskolin, N-acetyl-L-cysteine, R-spondin1 (Wnt agonists), FGF, EGF, and N2. Three different conditioned media were prepared by combining these niche factors in different ratios, as summarized in **Supplementary Table 2**.

### Hematoxylin & eosin (H&E), immunohistochemical (IHC), and immunofluorescence (IF) staining

Slides were prepared by cutting paraffin blocks to a thickness of 0.4 μm for H&E, IHC, and IF staining. First, one slide was subject to H&E staining after deparaffinization. For IHC staining, the following primary antibodies were used: p53 (CST, Cat# 2527S, 1:100), Ki-67 (CST, Cat# 9449S, 1:600), KRT5 (CST, Cat# 71536S, 1: 200), deltaNP63 (Abcam, Cat# ab203826, 1:100), p16 (Abcam, Cat#108349, 1:100), AREG (Invitrogen, Cat#MA5-41546, 1:100), phospho-AKT (CST, Cat#4060S, 1:100), phospho-ERK (CST, Cat#4370S, 1:100), Recombinant anti-human nucleoli (Abcam, Cat#90710, 1:200). Antigen retrieval was performed using Tris-EDTA (pH 9.0), and KRT5 was stained without antigen retrieval. For IF staining, primary antibodies including CD44 (CST, Cat# 3570, 1:100), ALDH1A1 (CST, Cat# 36671S, 1:400), E-cadherin (BD Transduction Laboratories, Cat# 610181, 1:50), and β-catenin (CST, Cat# 8814S, 1:400) were used and incubated overnight at 4°C. Secondary antibodies were next administered, including: donkey anti-rabbit IgG, Alexa Fluor™ Plus 488 (Invitrogen, Cat# A32790); donkey anti-mouse IgG, Alexa Fluor™ Plus 488 (Invitrogen, Cat# A32766); donkey anti-rabbit IgG, Alexa Fluor ™ Plus 594 (Invitrogen, Cat# A32754); and donkey anti-mouse IgG, Alexa Fluor™ Plus 594 (Invitrogen, Cat# A32744).

### Western blot assay

Harvested cells were lysed with RIPA buffer mixture containing cocktails of protease inhibitors (Santa Cruz Biotechnology, Cat# sc24948A) and mixed with Laemmli sample buffer (BioRad, Cat#161-0737). Proteins were loaded on SDS-PAGE gels (GenScript, Cat#M00669) and ran for 1 hour on 120 volts. Subsequently, the protein was transferred to 0.45μm PVDF membranes (Millipore Sigma, Cat#IPVH00010) with transfer buffer (formula: Mixture of 25mM Tris, 192mM Glycine, and 20% Methanol) for 2 hours on 270mA. Membranes were blocked by 2% BSA buffer diluted in 1X PBST (1x PBS with 1% Tween 20) for 1 hour at room temperature. The primary antibodies (anti-rabbit phospho-AKT, 1:1000; anti-rabbit phospho-ERK, 1:1000; anti-mouse AREG, 1:1000; anti-mouse ERK, 1: 1000; anti-rabbit AKT, 1:1000; anti-mouse β-actin, 1: 5000) in 2% BSA buffer were added to the membrane and incubated overnight at 4 °C. Membranes were then washed 3 times for 5 minutes with 1X PBST. The secondary antibody which is conjugated with HRP was added to the membrane and incubated for 1 hour at room temperature. Membrane was developed with ECL substrate (Thermo Fisher Scientific, Cat#A38554) and detected by chemiluminescence image reader (Fujifilm, LAS-4000). Broad multicolor pre-stained protein standard (Genescript, Cat#M00624-250) was used as a marker.

### RNA extraction, cDNA synthesis, and quantitative real-time PCR

Total RNA was extracted by the RNeasy Mini Kit (QIAGEN, Cat#70106). cDNA synthesis was performed using the Luna script RT Supermix kit (New England Biolabs, Cat#M3010L). Quantitative real-time PCR (qRT-PCR) was performed by using the Powerup™ SYBR Green Master Mix (Thermo Fisher Scientific, Cat#A25918). TATA Binding Protein (TBP) was used as the reference gene for normalization. The primers used in this study are listed in the **Supplementary Table 7**.

### Whole-exome sequencing (WES) and genomic analyses

Genomic DNA was extracted from tumor tissues, tumor organoids, and matched peripheral blood samples using an AllPrep DNA/RNA mini kit (Qiagen). DNA libraries were generated using the Agilent SureSelect Human All Exon V6 kit (Agilent Technologies). Sequencing was performed using the Illumina NovaSeq platform. Low-quality reads and adapters were removed using FastQC. GATK was used for the single-nucleotide variant (SNV) calling. Each read was mapped to the human reference genome (hg38) using the Burrows-Wheeler Alignment tool. Mutec2 was used for the SNV and indel analysis, and copy number alterations (CNAs) were analyzed using Sequenza, and CNVkit (*22, 23, 46*). The functional impact of mutations was predicted using the VEP method (*47*). To reduce false-positive variants, all variants were filtered using the following filtering criteria: (a) total depth < 50, (b) variant allelic frequency (VAF) < 7%, (c) altered read counts < 4. Further filtering criteria were applied to assess pathogenicity: (a) combined annotation dependent depletion (CADD) phred score < 26, and (b) minor allele frequency (MAF) > 0.1% in the gnomAD database.

### RNA sequencing and analysis

Total RNA was extracted from HNSCC organoids using the AllPrep DNA/RNA Mini Kit (Qiagen). Library preparation was performed using the NEBNext Ultra RNA libray Prep Kit for Illumina (NEB, USA). RNA sequencing was performed using the Illumina NovaSeq platform. Sequence reads were aligned to the hg38 genome using STAR. Low-quality and adapter reads were removed using FastQC. Further analysis was performed using DESeq2, and the HNSCC subtype enrichment score was calculated through the gene set variation analysis (GSVA) using the GSVA function.

### Survival analysis

In bulk tumor transcriptomes with available follow-up information, the survival analyses were performed by the R package “survminer” with Kaplan–Meier analyses. To determine the optimal cutpoint for this variable at once, we used the cutpoint function from the ’survminer’ R package.

### Single-cell RNA sequencing (scRNA-seq) Sample processing and data generation

HNSCC PDO samples were dissociated into single cells, and dead cells were removed using a dead cell removal kit (Miltenyi Biotec). Cell suspensions were processed using the Chromium Next GEM Single Cell 3ʹ Reagent Kits v.3.1 (10X Genomics) according to the manufacturer’s instructions. The resulting cDNA libraries All libraries were sequenced using the NovaSeq 6000 platform. BCL files were demultiplexed based on the 10X Genomics i7 index by using the bcl2fastq and mkfastq commands provided by 10X Genomics CellRanger v4.0.0.

### Data processing

Raw reads were aligned to the hg38 reference genome using Cell Ranger (v.7.0.1), and the expression matrices were generated per PDO. Raw data of the gene expression matrix was filtered using the "Seurat (v4.3.0)" R package with the following parameters: cells with > 200 genes and < 8000 genes, < 10% of mitochondrial gene expression in UMI counts, log10 (Genes Per UMI count) > 0.80, and genes expressed in > 0.1% cells. Doublets were removed using the R packages DoubletFinder (v2.0.3). After sequencing quality control and standardization, a total of 68,919 cells were obtained from 8 PDO lines (average=8,615 cells/PDO), with an average of 23,742 (22,202 ∼ 24,606) genes being measured in each cell.

The batch effects of the samples were corrected using the ‘RunHarmony’ function in the "harmony" (v0.1.1) R package. Dimension reduction was performed using UMAP with the ’RunUMAP’ function (dims = 1:50). Cell types were annotated using canonical markers, including KRT5, KRT14, EPCAM and SFN for squamous epithelial cells; COL1A2 and COL3A1 for fibroblasts; VWF and CDH5 for endothelial cells; CD19 and CD79A for B cells; CD3D and CD3E for T cells; LYZ and CLEC4C for myeloid cells.

### Identification of ITH program

To capture tumor ITH, we employed non-negative matrix factorization (NMF) on each of the 8 HNSCC PDOs, as described previously (*32*). Negative values in the centered expression matrix were adjusted to zero. NMF was performed individually for each PDO sample with a range of factors (K) from 6 to 9, resulting in 30 programs (or gene modules) per PDO. Each program was summarized by the top 50 genes based on NMF coefficients, yielding a total of 240 programs. To identify robust NMF programs that occurred across different K values and PDO lines, we applied three criteria. First, a program had to exhibit an overlap of at least 70% (35 of 50 genes) with a program with different K value within the same PDO. Second, a program shared an overlap of at least 20% (10 of 50 genes) with a program in any other PDO line. The NMF programs were ranked based on their similarity with programs from other PDOs, and selection was performed in a decreasing order. Third, once a program was chosen, any other program within the same PDO that had an overlap of 20% or more with the selected NMF program was excluded to reduce redundancy. This process resulted in 50 final NMF programs, which were then subjected to hierarchical clustering using the unweighted pair group method with the arithmetic mean method based on Jaccard similarity. As a result, eight clusters of consensus ITH programs were identified. For each consensus ITH program, genes that occurred in at least 40% of the constituent programs were defined as program signature genes.

To understand these eight consensus ITH programs, we performed comparison analysis against 41 curated gene programs (termed meta-program or MP in the original study) established by the same NMF approach from scRNA-seq delineation of pan-cancer tumor samples (*15*). First, we calculated the average similarity (Jaccard index) and single-cell score correlation between each program in MP and each in our consensus ITH programs. We assessed statistical significance by permuting our ITH programs 100 times and considered a confidence threshold of 99.9% as significant. In addition, we collected scRNA-seq-defined program signature genesets for three common types of SCCs, namely HNSCC, esophageal SCC (ESCC), cervical SCC (CSCC) (*26, 31, 48*) and validated our annotations using Jaccard index and hypergeometric tests.

### Scoring the ITH programs in each single cell using a random geneset approach

For each of the 8 consensus ITH programs, 1000 background genesets with comparable expression levels were randomly generated as described previously (*49*). For each single cell, the average centered expression of ITH program geneset as well as the 1000 randomly-generated genesets was measured. P value was defined as the proportion of randomly-generated genesets which were expressed higher than the ITH program geneset. The ITH program score was then defined as −log10(p) and rescaled linearly to [0,1].

This method was also applied to determine the molecular subtype enrichment score of each single cell using the published genesets for each molecular subtype, as an independent method to validate the GSVA approach (*24*).

To directly compare our PDO single cell ITH program scores with *in vivo* HNSCC primary tumor single cell data, we selected HNSCC PDOs or tumors harboring the hEMT and EpiSen programs and combined our PDOs and primary tumors into joint datasets (*14*). Expression levels (log2 (TPM+1)) of each individual dataset were mean centered per gene before the cells were combined. Principal component analysis (PCA) was then performed on the joint datasets using the top 4,500 genes with the highest expression levels. For visualization, we subsequently selected the top 2 principal components (PCs) that exhibited a strong correlation (r>0.3 or r<−0.3) with the single-cell scores corresponding to ITH programs of interest.

### Animal and xenograft experiments

Animal experiments were performed according to the guidelines for animal experiments established by the Yonsei University Institutional Animal Care and Use Committee. Five-week-old female BALB/c nude mice were purchased from the Shizuoka Laboratory Animal Center (Shizuoka, Japan) and reared in a specific pathogen-free environment. Prior to the experiment, the animals were acclimated for 1 week and had free access to food and water. PDO lines were first dissociated into single cells, and 2.5 x 10^5^ cells were mixed in 100 μl of Matrigel and injected subcutaneously into both flanks of each animal (n=3/group). Subsequently, changes in tumor size were measured continually. The tumor volume was calculated using the formula (volume = (length x width^2^) / 2).

### Data acquisition

Bulk HNSCC RNA-seq and proteomics datasets were collected from The Cancer Genome Atlas (TCGA, RRID:SCR_003193), and Clinical Proteomic Tumor Analysis Consortium (CPTAC, RRID:SCR_017135), as well as The Human Protein Atlas (HPA, RRID:SCR_006710). (*24, 33, 35*) HNSCC cell line RNA-seq data were obtained from the The Cancer Cell Line Encyclopedia (CCLE) dataset (*36*).

### Statistical analysis

All statistical analyses were conducted on R (Version 3.5.0). Wilcoxon rank-sum tests were used for comparing noncategorical values between groups. The x2 test was used to compare associations between categorical variables. Pearson’s correlation was used to compare the similarity among subgroups. For survival analysis, log-rank test was used for statistical analyses among Kaplan–Meier curves. Two-tailed P values < 0.05 was used for all other statistical significance unless stated elsewhere.

## Acknowledgment

This work was supported by National Research Foundation of Korea (NRF) grant funded by the Korean government (MSIT) (NRF-2021R1C1C1012004) (Y.M.P.)

NIH under the award R37CA237022 and 1R01DE033648 (D.-C.L.)

Ming Hsieh Institute for Research of Engineering-Medicine for Cancer (D.-C.L.).

## Author contributions

Conceive and design study: Y.M.P. and D.-C.L. Design experiments and analysis: Y.M.P. and D.-C.L.

Perform the experiments: J.H.U., C.N., H.Z., and S.-J.S.

Perform bioinformatics and statistical analysis: Y.Z., Q.M., and Y.M.P

Contribute the reagents and materials: Y.W.K. and S.-J.S.

Analyze the data: J.H.U., Y.Z., Q.M., and C.N.

Supervise the research: Y.M.P. and D.-C.L.

Write the manuscript: J.H.U., C.N., Y.M.P., and D.-C.L.

## Data availability

The sequencing data generated in this study have been deposited in the SRA (PRJNA960652). All other relevant data supporting the key findings of this study are available within the article or from the corresponding author upon reasonable request.

## List of Supplementary Materials

Supplementary Figures: Supplementary figure 1 to 8 with figure legends

Supplementary tables: Table 1 to 7

Reference: 1 to 49

---

**Supplementary Figure 1.**
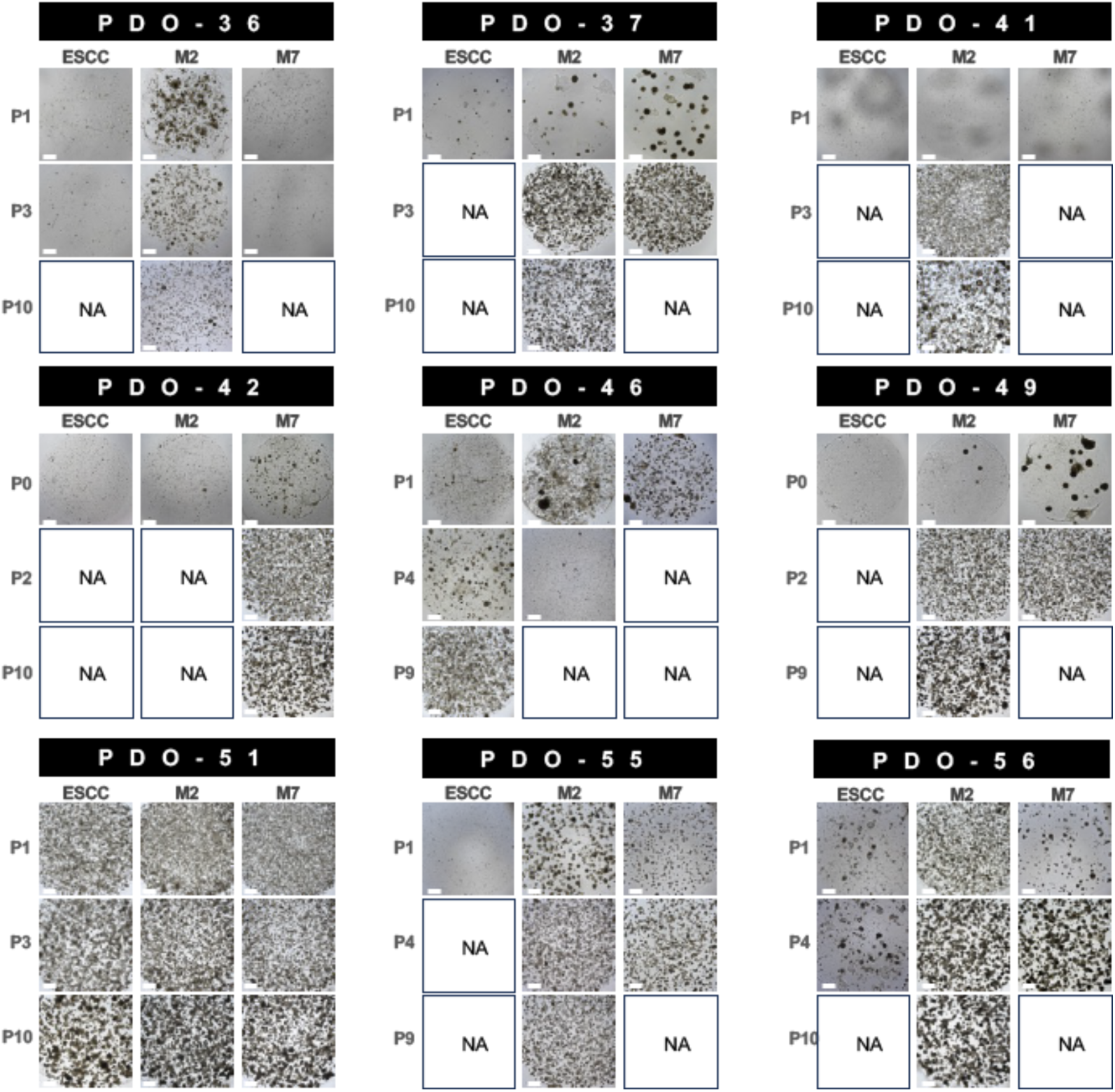
Images of exemplary PDOs grown in different culture media across different time points.

**Supplementary Figure 2.**
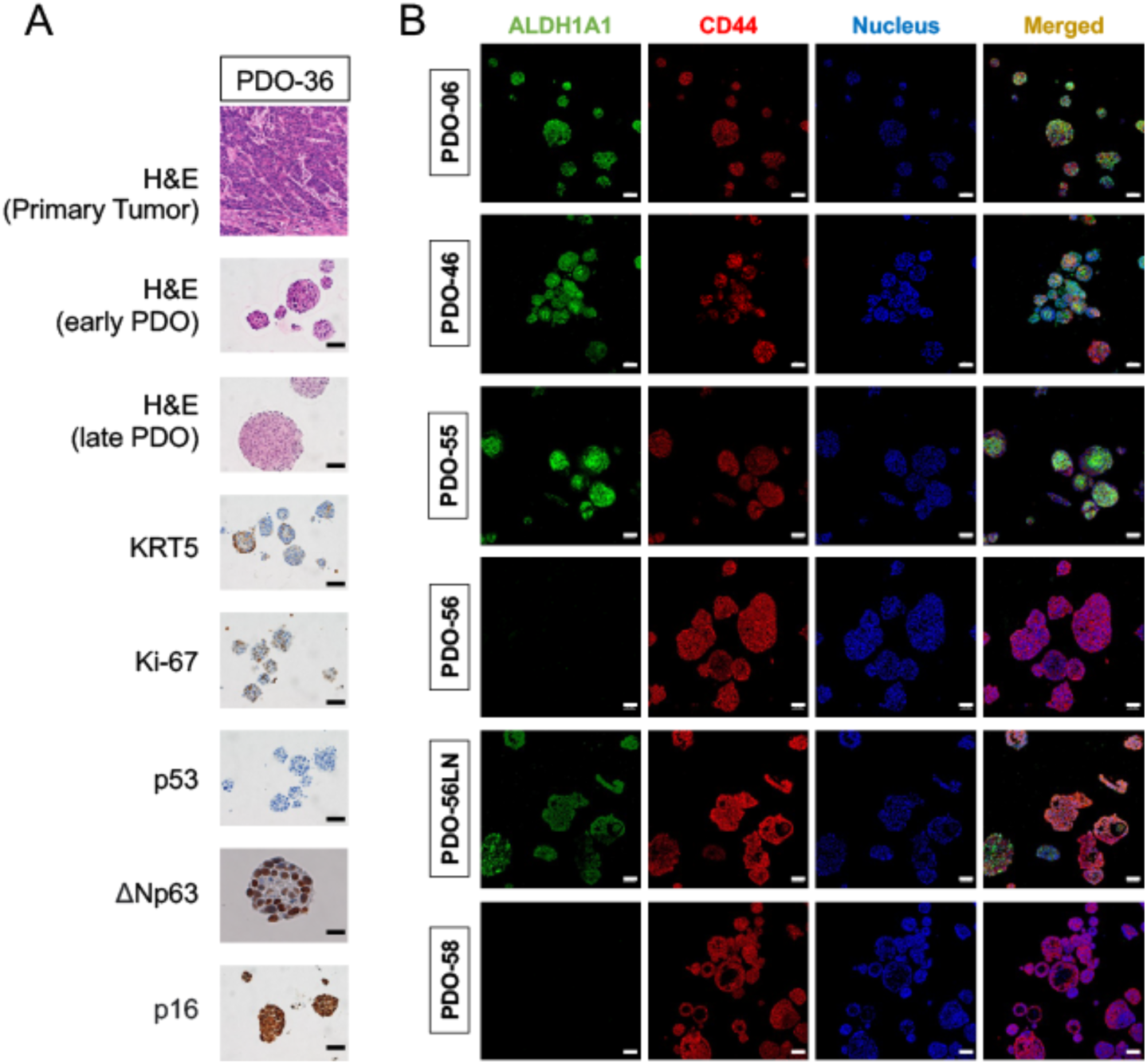
(**A**) H&E and IHC staining of indicated markers on PDO-36 (HPV+) and its matched primary tumor. (**D)** The expression of stem cells markers (ALDH1A1 and CD44) was evaluated using IF staining in selected PDO lines.

**Supplementary Figure 3.**
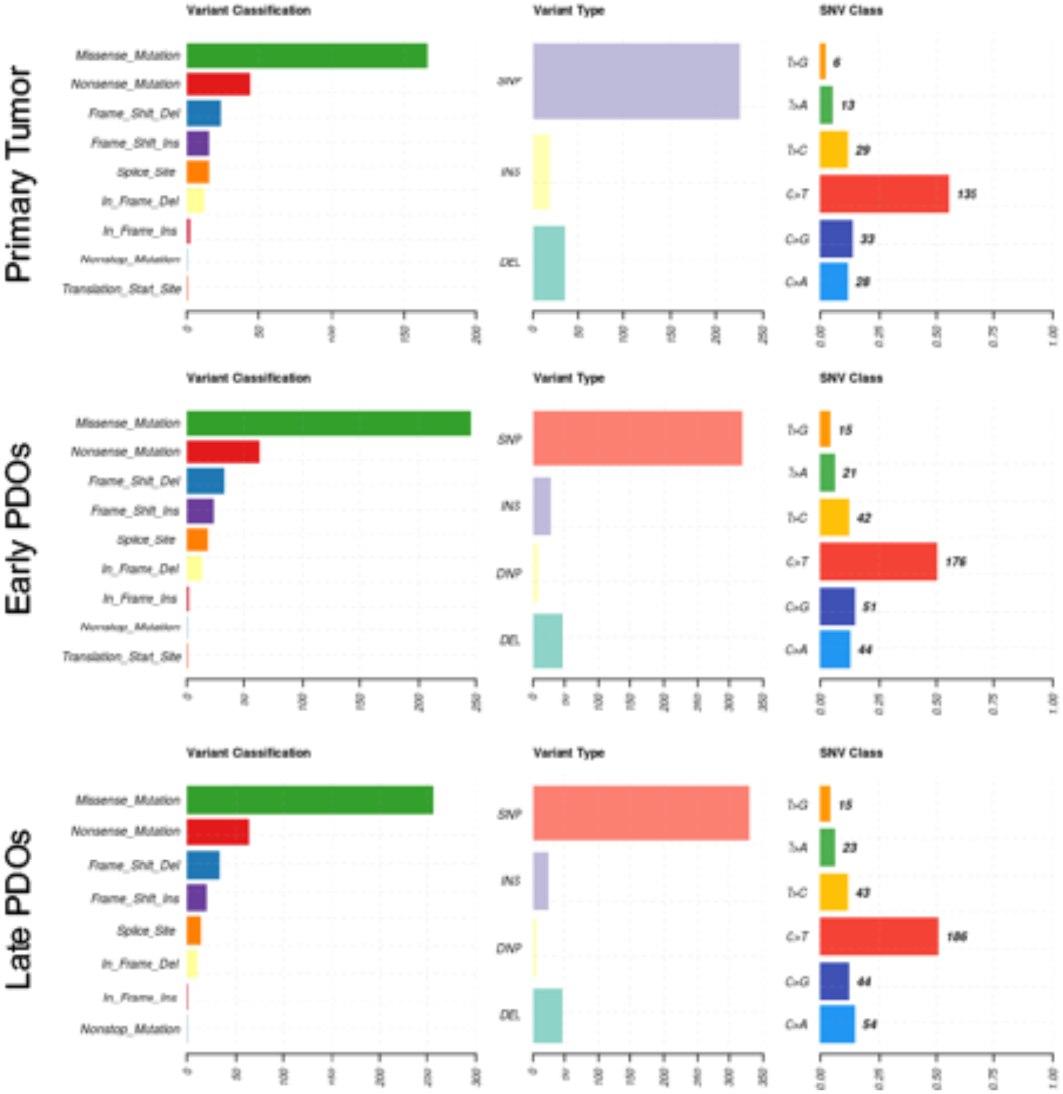
Somatic variant characteristics of primary tumors, early- and late-passaged PDOs.

**Supplementary Figure 4.**
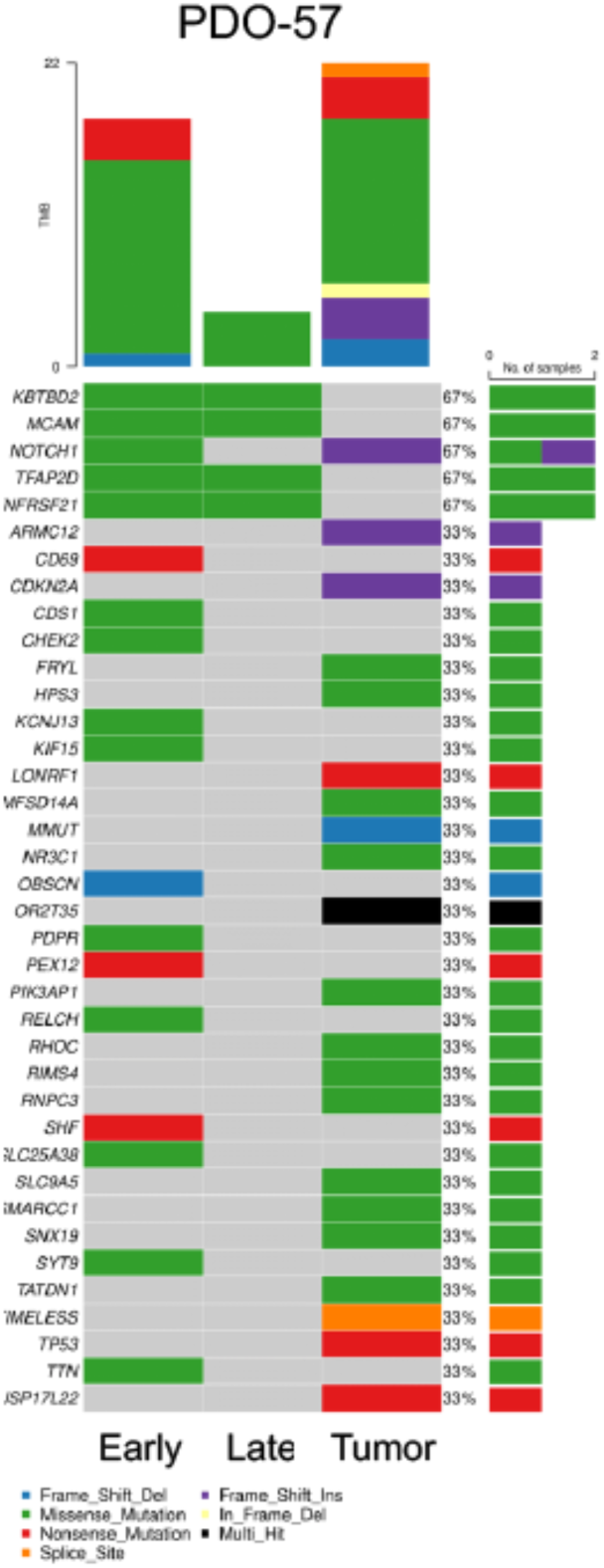
Oncoplot comparing somatic mutations between the primary tumor, early- and late-passaged PDO-57.

**Supplementary Figure 5.**
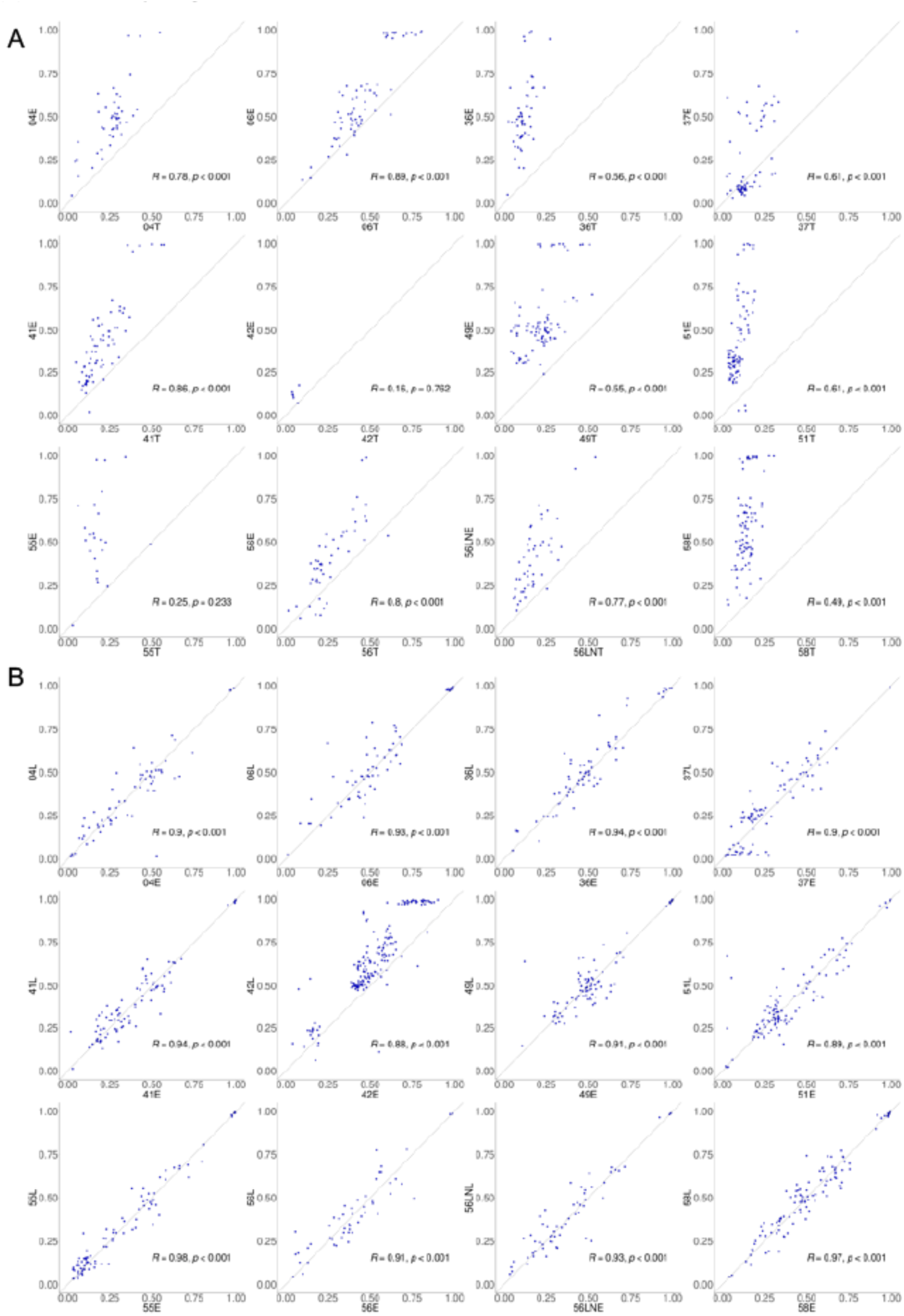
**(A)** Scatter plots comparing mutational VAFs between primary tumor (T) vs. matched early-passaged (E) PDOs (E), and (**B**) between early- and late-passaged (L) PDOs.

**Supplementary Figure 6.**
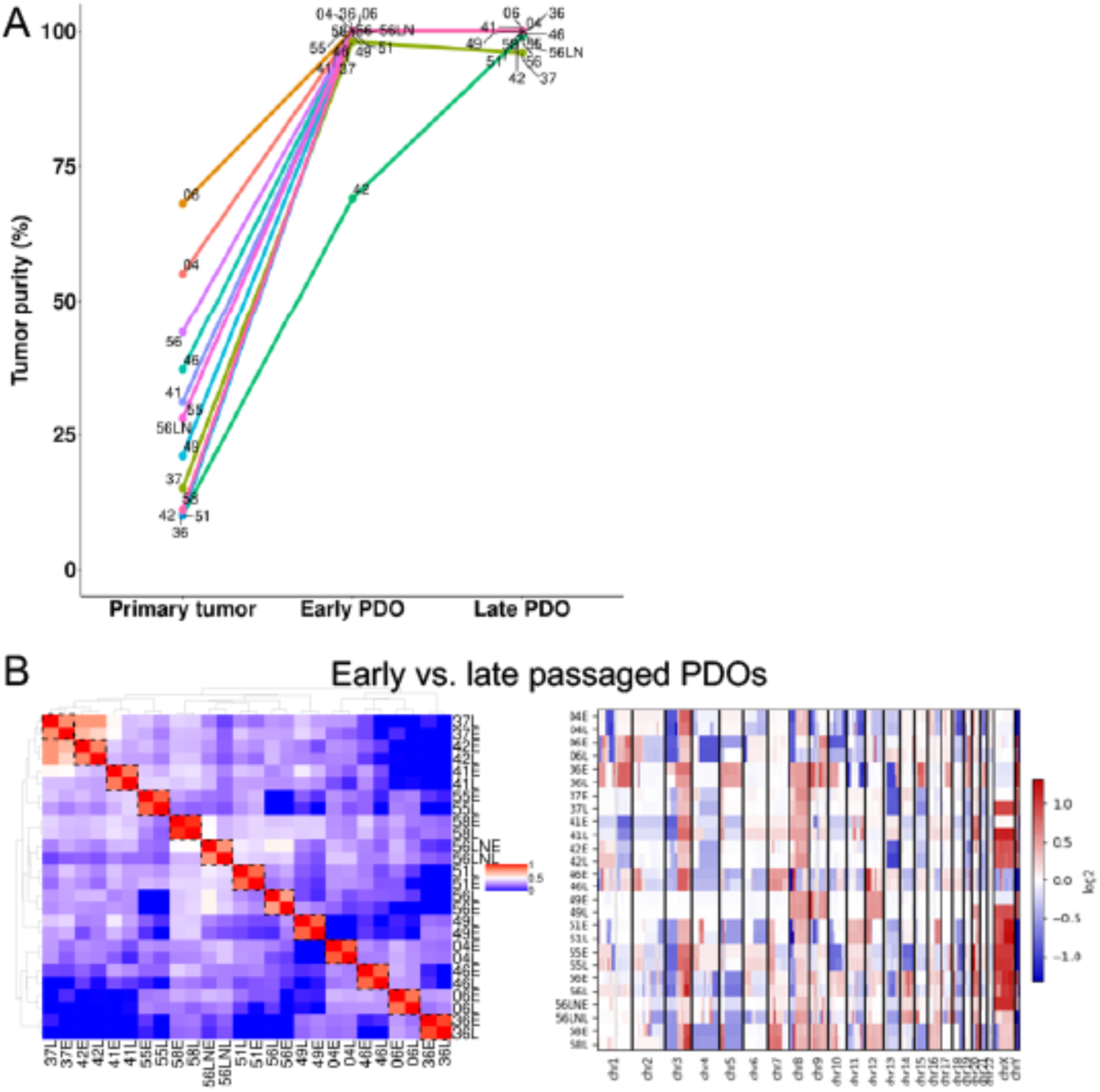
**(A)** Line plots showing the tumor cellularity of each matched primary tumor, early- and late-passaged PDOs. **(B)** Left, heatmap showing the unsupervised clustering of early- and late-passaged PDOs. Right, CNV profiles of early- and late-passaged PDOs.

**Supplementary Figure 7.**
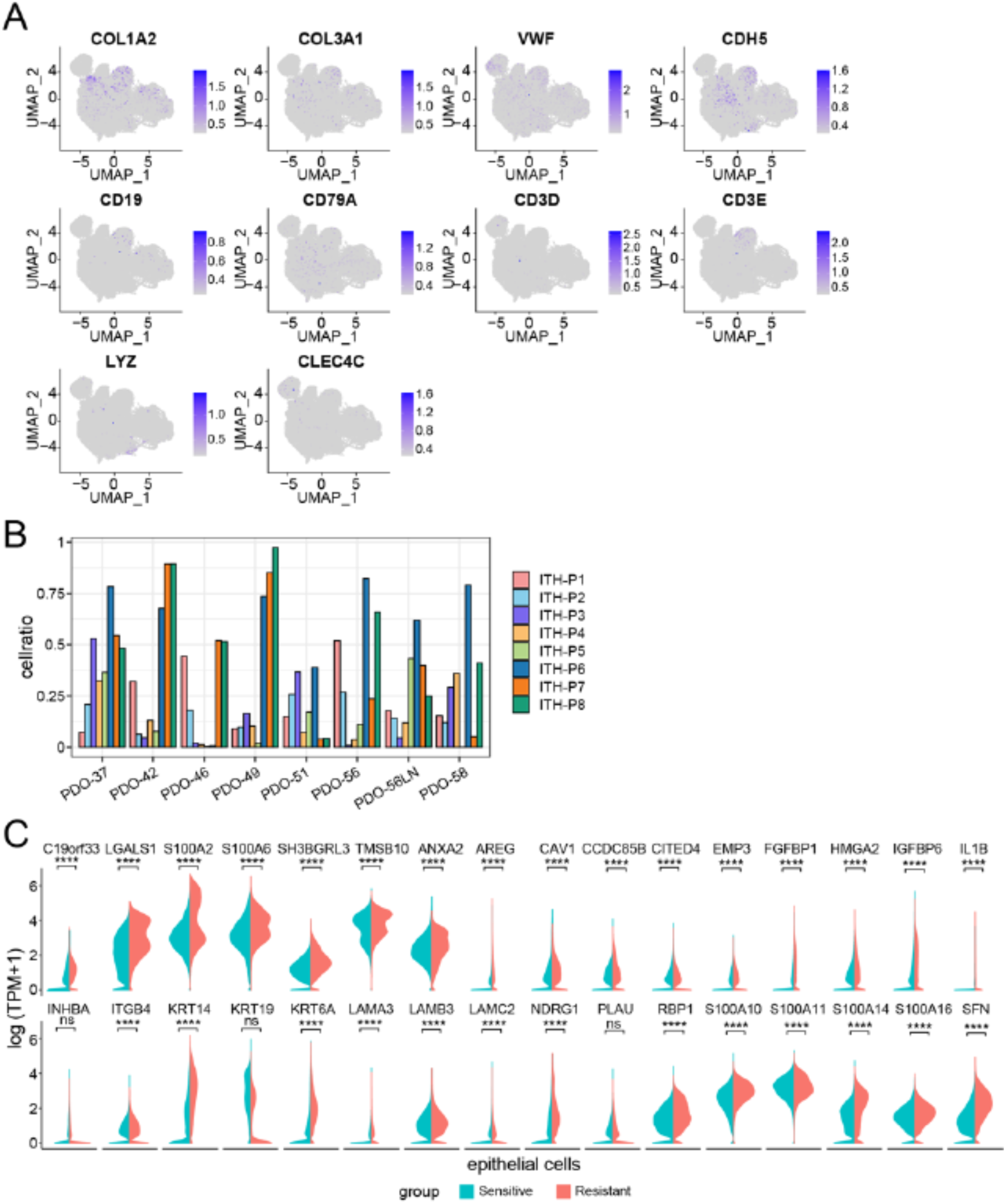
**(A)** UMAP of canonical markers for various stromal cell types, including COL1A2 and COL3A1 for fibroblasts; VWF and CDH5 for endothelial cells; CD19 and CD79A for B cells; CD3D and CD3E for T cells; LYZ and CLEC4C for myeloid cells. **(B)** Column plots showing the cell ratio of each ITH program in each PDO line. **(C)** Violin plots showing the expression of hEMT genes in cisplatin resistant vs. sensitive PDO samples.

**Supplementary Figure 8.**
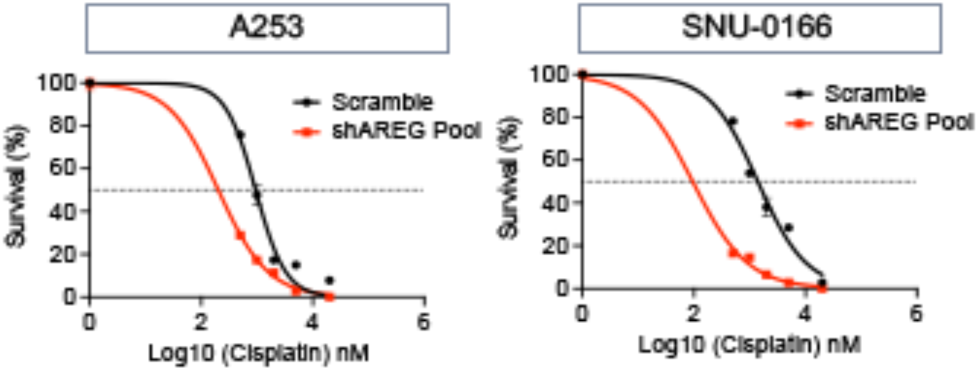
Dose-response curves of scramble control and shAREG-knockdown samples to cisplatin treatment in HNSCC cell lines (A253 and SNU-1066).

