## Supplementary Figure for "Genomic and single-cell characterization of patient-derived tumor organoid models of head and neck squamous cell carcinoma"

This document contains

8 supplementary figures and legends: S1 to S8

#### Supplementary Figure Legends

**Supplementary Figure 1.** Images of exemplary PDOs grown in different culture media across different time points.

**Supplementary Figure 2.** (**A**) H&E and IHC staining of indicated markers on PDO-36 (HPV+) and its matched primary tumor. (**D)** The expression of stem cells markers (ALDH1A1 and CD44) was evaluated using IF staining in selected PDO lines.

**Supplementary Figure 3.** Somatic variant characteristics of primary tumors, early- and late-passaged PDOs.

**Supplementary Figure 4.** Oncoplot comparing somatic mutations between the primary tumor, early- and late-passaged PDO-57.

**Supplementary Figure 5. (A)** Scatter plots comparing mutational VAFs between primary tumor (T) vs. matched early-passaged (E) PDOs (E), and (**B**) between early- and late-passaged (L) PDOs.

**Supplementary Figure 6. (A)** Line plots showing the tumor cellularity of each matched primary tumor, early- and late-passaged PDOs. **(B)** Left, heatmap showing the unsupervised clustering of early- and late-passaged PDOs. Right, CNV profiles of early- and late-passaged PDOs.

**Supplementary Figure 7. (A)** UMAP of canonical markers for various stromal cell types, including COL1A2 and COL3A1 for fibroblasts; VWF and CDH5 for endothelial cells; CD19 and CD79A for B cells; CD3D and CD3E for T cells; LYZ and CLEC4C for myeloid cells. **(B)** Column plots showing the cell ratio of each ITH program in each PDO line. **(C)** Violin plots showing the expression of hEMT genes in cisplatin resistant vs. sensitive PDO samples.

**Supplementary Figure 8.** Dose-response curves of scramble control and shAREG-knockdown samples to cisplatin treatment in HNSCC cell lines (A253 and SNU-1066).

### Supplementary Figure 1

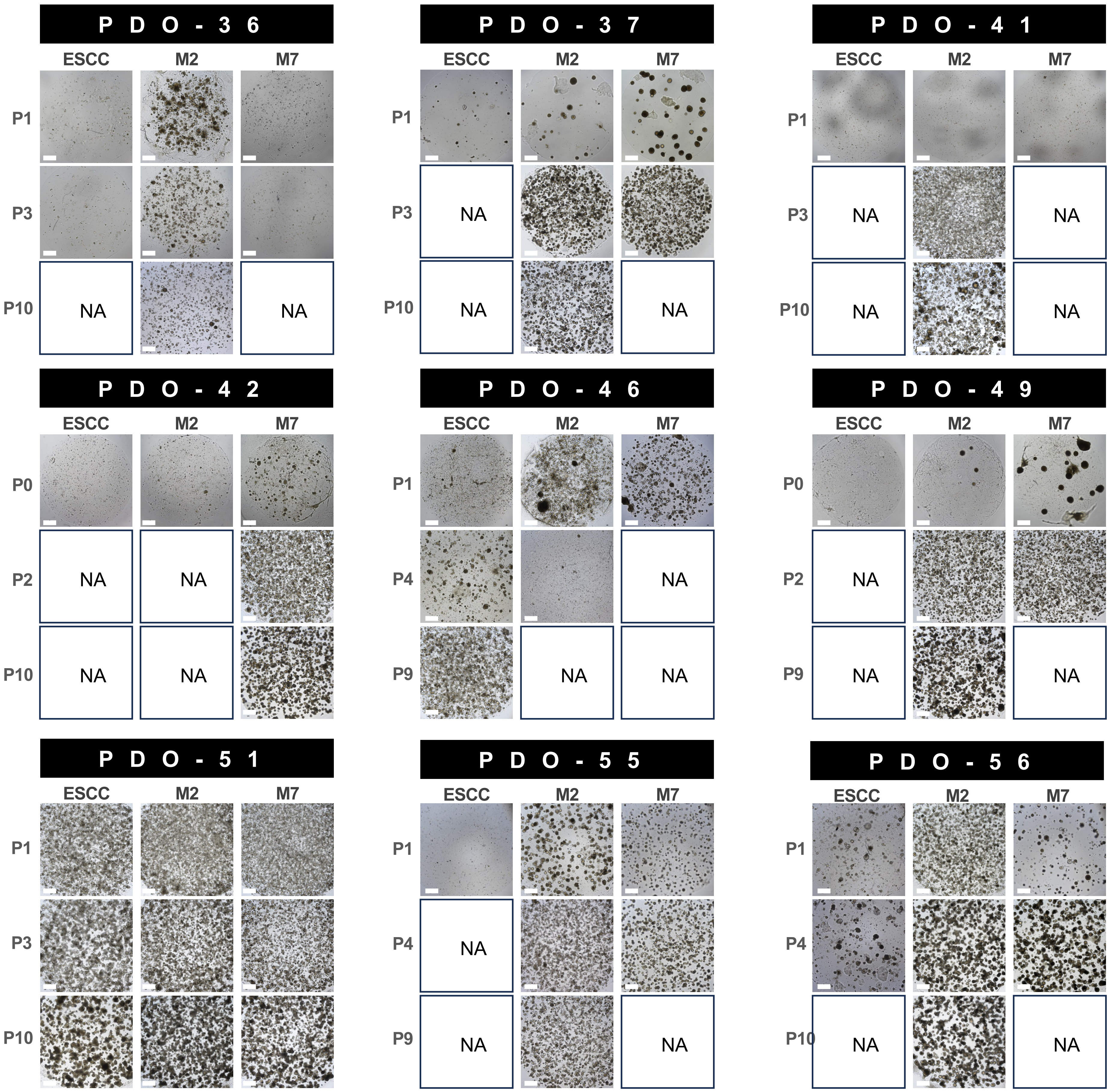

### Supplementary Figure 2

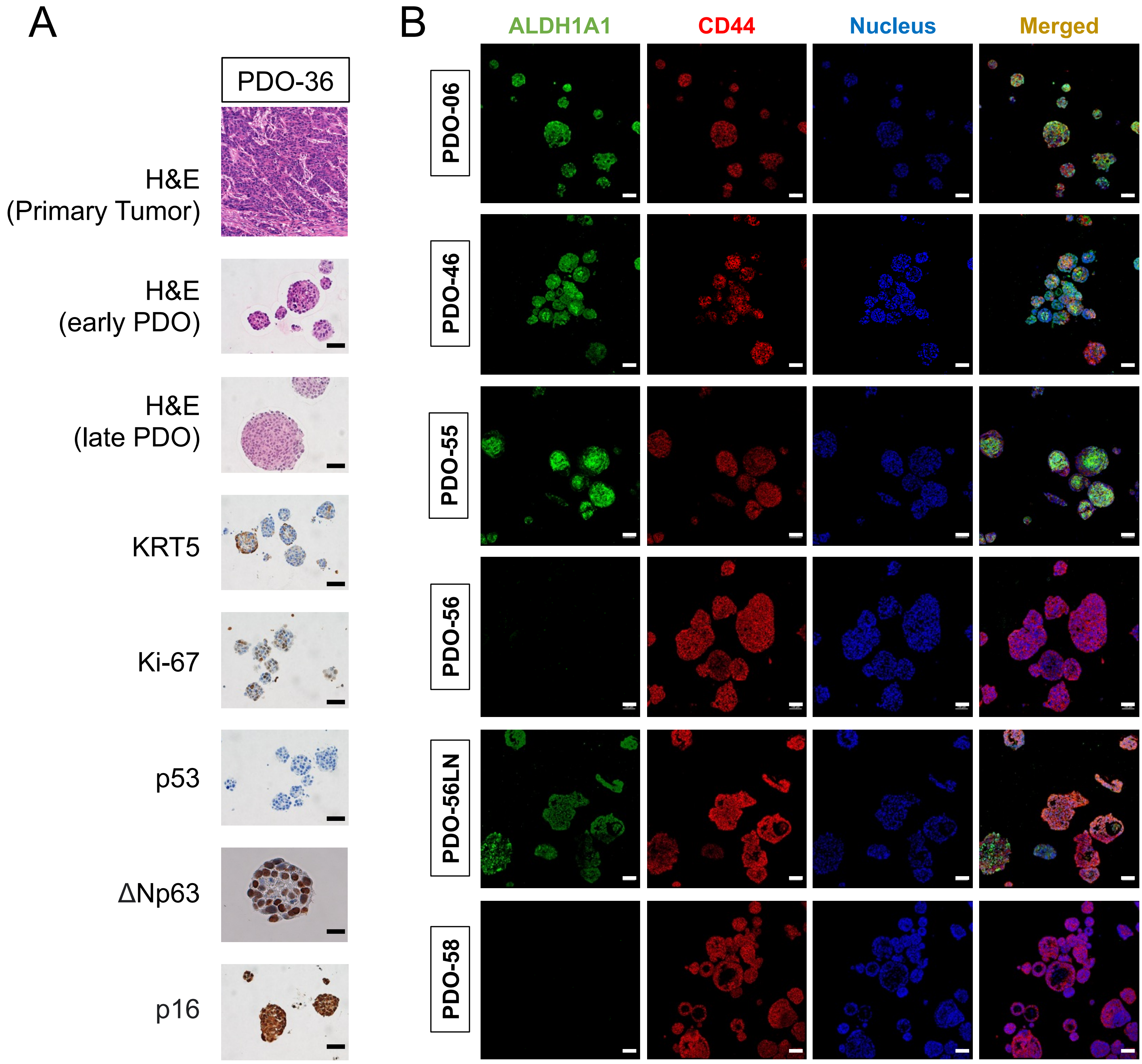

Supplementary Figure 3

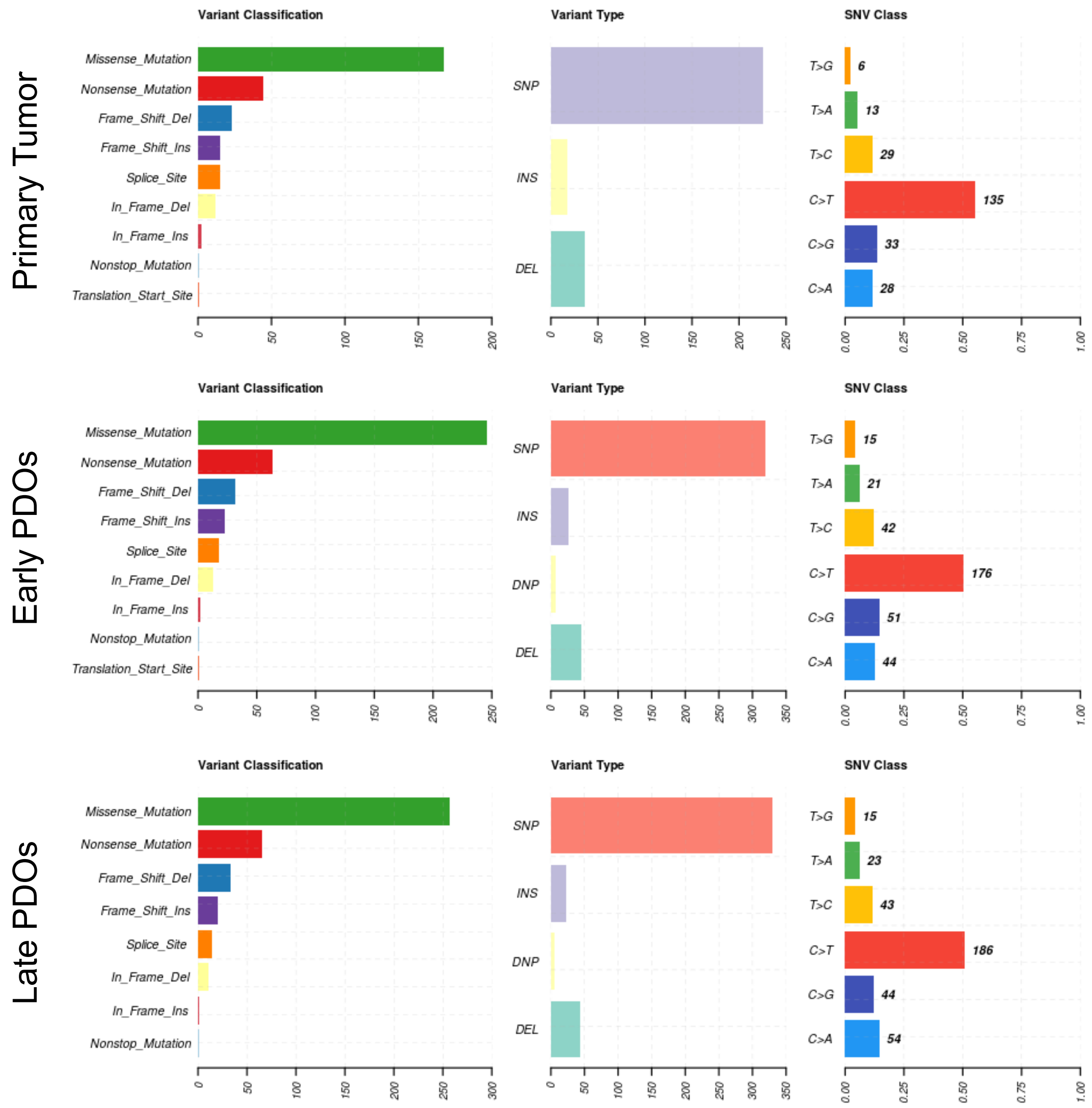

### Supplementary Figure 4

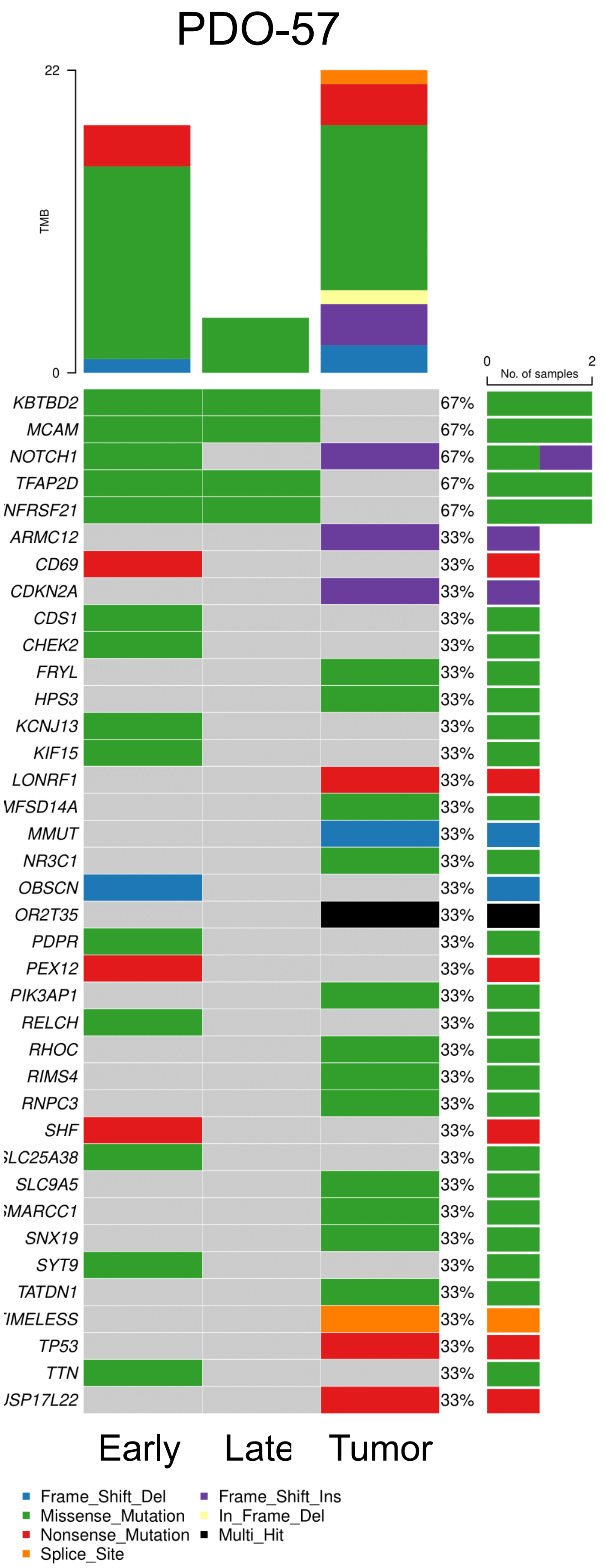

### Supplementary Figure 5

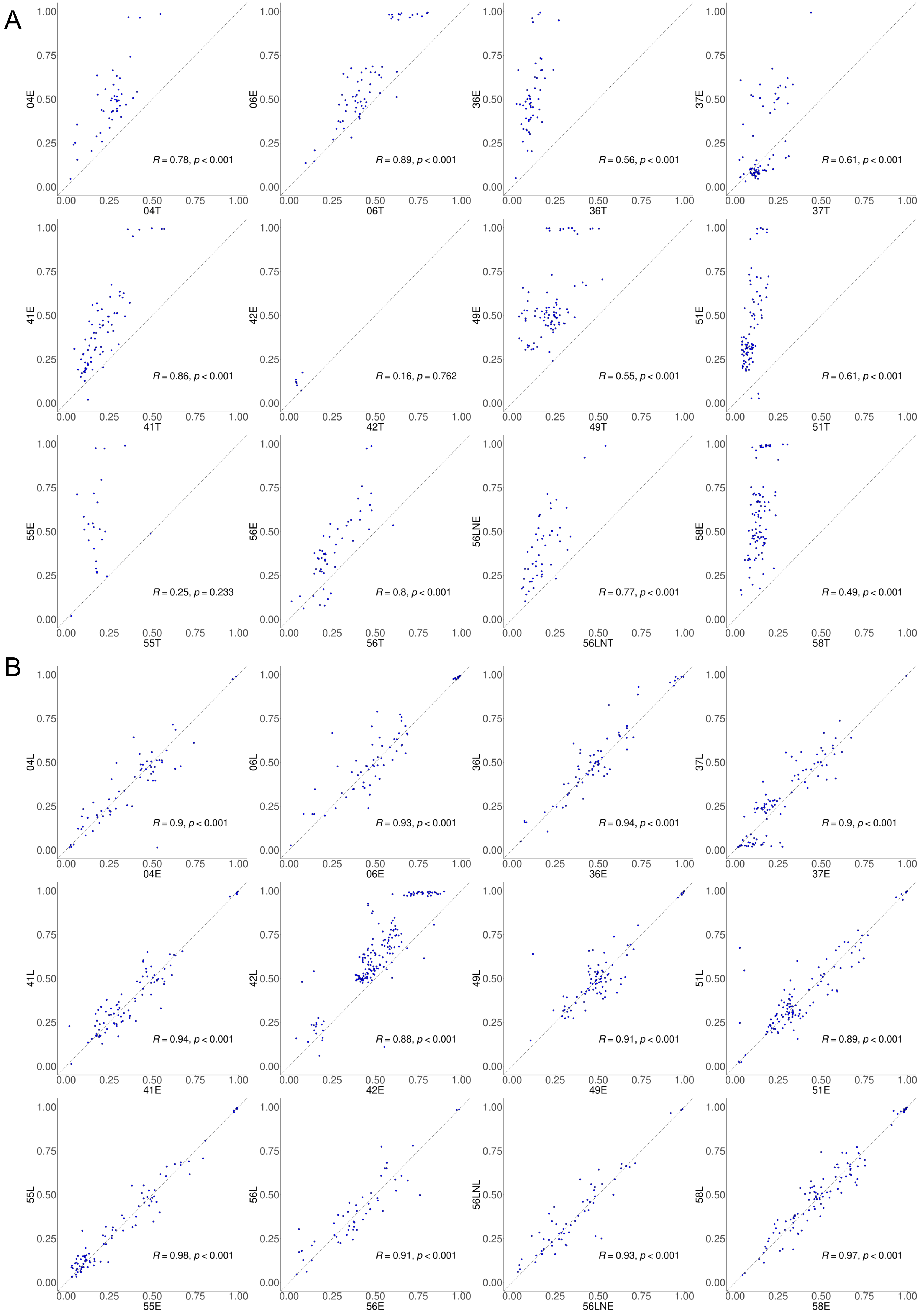

### Supplementary Figure 6

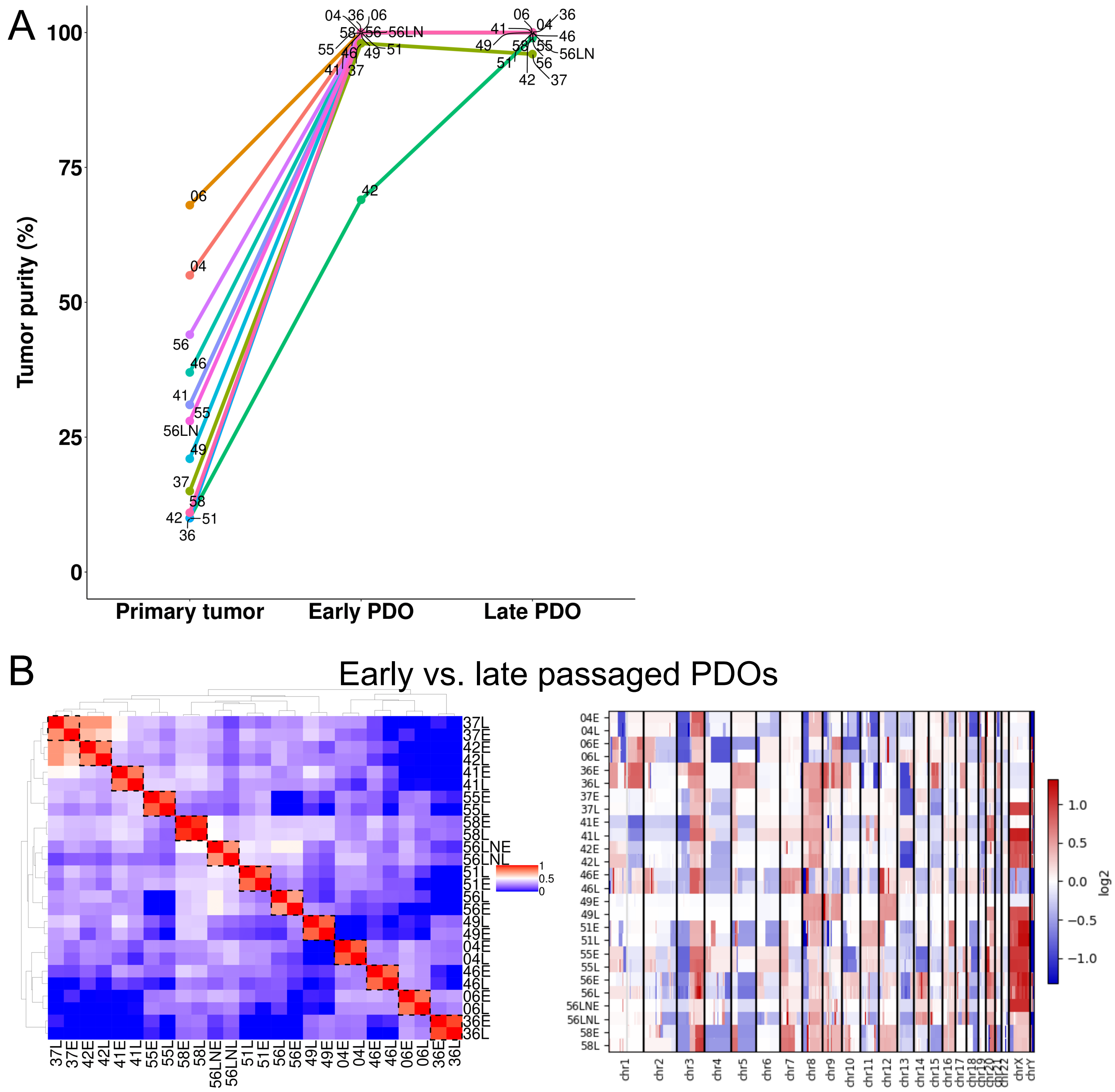

### Supplementary Figure 7

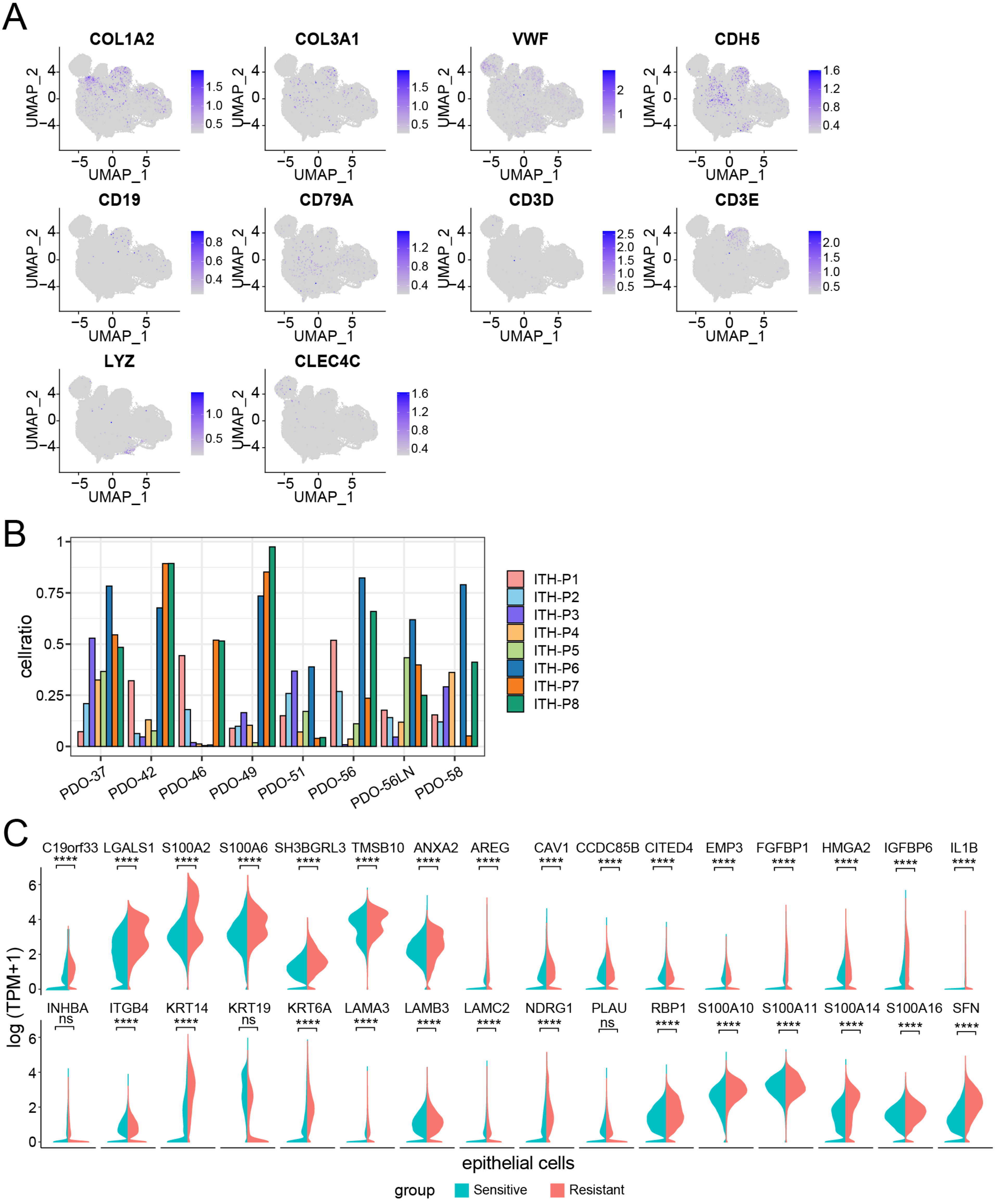

### Supplementary Figure 8

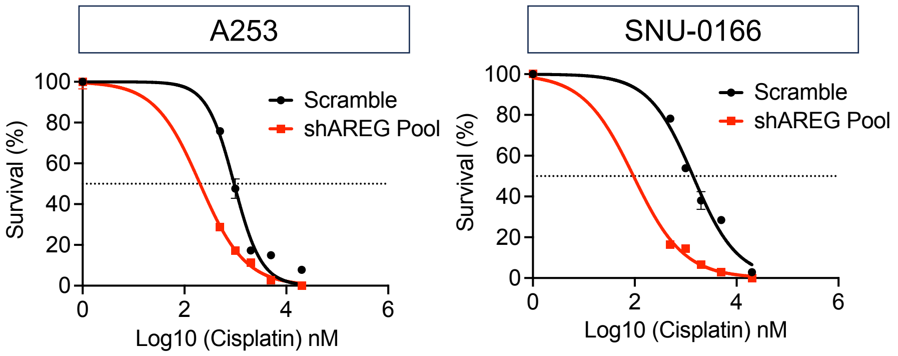
