## Supplementary Table for "Genomic and single-cell characterization of patient-derived tumor organoid models of head and neck squamous cell carcinoma"

**Supplementary Table 1.** Clinico-pathological data of 31 patients in this study

| **No.** | **PDO ID** | **Age** | **Sex** | **Site** | **Path** | **p16** | **pT** | **pN** | **LVI** | **PNI** | **ENE** |
| --- | --- | --- | --- | --- | --- | --- | --- | --- | --- | --- | --- |
| 1 | 04 | 35 | M | OC | SCCa | - | 3 | 1 | Yes | Yes | No |
| 2 | 06 | 65 | M | HPx | SCCa | - | 4a | 2a | Yes | Yes | Yes |
| 3 | 36 | 65 | M | OPx | SCCa | + | 2 | 1 | Yes | Yes | Yes |
| 4 | 37 | 47 | M | OC | SCCa | - | 3 | 3b | No | Yes | Yes |
| 5 | 41 | 41 | M | OPx | SCCa | - | 2 | 0 | No | No | No |
| 6 | 42 | 23 | M | OC | SCCa | - | 2 | 2a | Yes | Yes | Yes |
| 7 | 46 | 60 | M | HPx | SCCa | - | 2 | 3b | Yes | No | Yes |
| 8 | 49 | 65 | M | OC | SCCa | - | 3 | 0 | No | No | No |
| 9 | 51 | 46 | F | OC | SCCa | - | 2 | 0 | No | Yes | No |
| 10 | 55 | 57 | F | OC | SCCa | - | 4a | 1 | No | No | No |
| 11 | 56 | 55 | M | OC | SCCa | - | 3 | 3b | No | No | Yes |
| 12 | 57 | 40 | M | OC | SCCa | - | 2 | 3b | Yes | No | Yes |
| 13 | 58 | 61 | M | HPx | SCCa | - | 4a | 2b | Yes | No | No |
| 14 | 63 | 51 | M | OPx | SCCa | - | 3 | 0 | No | No | No |
| 15 | 65 | 48 | F | OPx | SCCa | - | 2 | 0 | No | No | No |
| 16 | 66 | 55 | M | OC | SCCa | - | 2 | 1 | No | Yes | Yes |
| 17 | 67 | 75 | M | OC | SCCa | - | 2 | 0 | No | No | No |
| 18 | 68 | 60 | F | OC | SCCa | - | 3 | 0 | No | No | No |
| 19 | 69 | 65 | M | OPx | SCCa | - | 1 | 2b | No | No | Yes |
| 20 | 70 | 53 | M | OC | SCCa | - | 4 | 3b | No | No | Yes |
| 21 | 71 | 69 | M | Lx | SCCa | - | 2 | 2a | No | No | Yes |
| 22 | failed | 61 | M | OPx | SCCa | + | 2 | 2a | No | No | Yes |
| 23 | failed | 31 | M | OC | SCCa | - | 2 | 2b | Yes | Yes | No |
| 24 | failed | 67 | M | OC | SCCa | - | 4 | 0 | No | No | No |
| 25 | failed | 62 | M | HPx | SCCa | - | 3 | 2b | Yes | Yes | No |
| 26 | failed | 83 | F | OC | SCCa | - | 3 | 2a | No | No | Yes |
| 27 | failed | 56 | F | OC | SCCa | - | 2 | 2a | No | Yes | No |
| 28 | failed | 60 | F | OC | SCCa | - | 2 | 2b | Yes | No | Yes |
| 29 | failed | 83 | F | OC | SCCa | - | 1 | 2b | Yes | No | Yes |
| 30 | failed | 49 | M | OC | SCCa | - | 3 | 1 | Yes | No | No |
| 31 | failed | 65 | M | Lx | SCCa | - | 2 | 2a | Yes | No | Yes |

No: number, PDOs: patient-derived tumor organoids, Path: pathology, OC: oral cavity, HPx: hypopharynx, OPx: oropharynx, Lx: larynx, SCCa: squamous cell carcinoma, LVI: lymphovascular invasion, PNI: perineural invasion, ENE: extranodal extension. p16 protein staining was performed using IHC.

**Supplementary Table 2.** Conditioned medium used for culturing and maintaining PDOs

|  | Stock con. | A | B | C |
| --- | --- | --- | --- | --- |
| Glutamax | 100 x | 1 x | 1 x | 1 x |
| HEPES | 100 x | 1 x | 1 x | 1 x |
| PS | 100 x | 1 x | 1 x | 1 x |
| Primocin | 500 x | 1 x | 1 x | 1 x |
| B27 (+Vit A) | 50 x | 1 x | 1 x | 1 x |
| Noggin | CM | 2 % | 1 % | 1 % |
| Y27632 | 100 mM | 10 μM | 10 μM | 10 μM |
| NIC | 2 M | - | 2.5 mM | 2.5 mM |
| SB202190 | 20 mM | - | 1 μM | 1 μM |
| A83-01 | 50 mM | - | - | 500 nM |
| Forskolin | 20 mM | - | - | 10 μM |
| FGF10 | 250 ng/μl | - | - | 100 ng/ml |
| FGF7 | 100 ng/μl | - | 25 ng/ml | 25 ng/ml |
| n-Acetyl cysteine | 500 mM | 1 mM | - | 1.25 mM |
| RSPO1 | CM | 2 % | - | 10 % |
| hEGF | 100 ng/μl | 50 ng/ml | - | - |
| N2 | 100 x | 1 x | - | - |

con: concentration, PS: penicillin-streptomycin, NIC: nicotinamide, RSPO1: R-spondin 1.

**Supplementary Table 3.** Optimal culture medium of 22 PDO lines

| No | PDO ID | Sex | Age | Sample site | Optimal medium |
| --- | --- | --- | --- | --- | --- |
| 1 | 04 | M | 35 | Primary tumor | A |
| 2 | 06 | M | 65 | Primary tumor | A |
| 3 | 36 | M | 65 | Primary tumor | B |
| 4 | 37 | M | 47 | Primary tumor | B |
| 5 | 41 | M | 41 | Primary tumor | C |
| 6 | 42 | M | 23 | Primary tumor | C |
| 7 | 46 | M | 60 | Primary tumor | A |
| 8 | 49 | M | 65 | Primary tumor | B |
| 9 | 51 | F | 46 | Primary tumor | A, B, C |
| 10 | 55 | F | 57 | Primary tumor | B, C |
| 11 | 56 | M | 55 | Primary tumor | A, B, C |
| 12 | 56LN | M | 55 | Metastatic LN | A, B, C |
| 13 | 57 | M | 40 | Primary tumor | C |
| 14 | 58 | M | 61 | Primary tumor | B, C |
| 15 | 63 | M | 51 | Primary tumor | C |
| 16 | 65 | F | 48 | Primary tumor | C |
| 17 | 66 | M | 55 | Primary tumor | B |
| 18 | 67 | M | 75 | Primary tumor | A |
| 19 | 70 | M | 53 | Primary tumor | B |
| 20 | 72 | M | 38 | Primary tumor | B |
| 21 | 73 | M | 36 | Primary tumor | B |
| 22 | 75 | M | 72 | Primary tumor | C |

PDOs: patient tumor derived organoids.

**Supplementary Table 4.** Correlation of VAF between primary tumor, early and late PDO

| PDO ID | Tumor *vs.* early PDO | Early *vs*. late PDO |
| --- | --- | --- |
| 04 | 0.79 | 0.90 |
| 06 | 0.88 | 0.92 |
| 36 | 0.56 | 0.94 |
| 37 | 0.61 | 0.90 |
| 41 | 0.85 | 0.94 |
| 42 | 0.16 | 0.89 |
| 46 | 0.71 | 0.97 |
| 49 | 0.53 | 0.91 |
| 51 | 0.60 | 0.88 |
| 55 | 0.24 | 0.97 |
| 56 | 0.80 | 0.90 |
| 56LN | 0.75 | 0.92 |
| 58 | 0.49 | 0.96 |

**Supplementary Table 5.** Signature genes of 8 ITH programs

| ITH-P1 | ITH-P2 | ITH-P3 | ITH-P4 | ITH-P5 | ITH-P6 | ITH-P7 | ITH-P8 |
| --- | --- | --- | --- | --- | --- | --- | --- |
| HSPE1 | BIRC5 | MAML2 | FTX | AGR2 | BNIP3 | KRT16 | C19orf33 |
| ATP5MC1 | CDK1 | RBMS3 | LINC01876 | LCN2 | DDIT4 | KRT17 | LGALS1 |
| C1QBP | CDKN3 | RUNX1 | MALAT1 | SAT1 | GAPDH | KRT6A | S100A2 |
| RAN | CKS1B | SOX6 | NEAT1 | SLPI | CRIP1 | KRT6B | S100A6 |
| SNRPD1 | CKS2 | AC013652.1 | ZFPM2-AS1 | CP | FAM162A | PERP | SH3BGRL3 |
| CYCS | HIST1H4C | ARL15 | ATXN1 | CXCL1 | VIM | RHCG | TMSB10 |
| H2AFZ | HMGB2 | DPYD | EXT1 | CXCL8 | C4orf3 | S100A9 | ANXA2 |
| HSPA8 | NUSAP1 | FBXO32 | LRMDA | EHF | CA9 | SERPINB3 | AREG |
| LDHB | STMN1 | HIVEP3 | PTK2 | RHOV | CLEC2B | TACSTD2 | CAV1 |
| NME1 | TOP2A | KANK1 | RAD51B | WFDC2 | IER3 | ANXA1 | CCDC85B |
| NME2 | TUBA1B | NFIB | TANC2 | ALCAM | KRT15 | CSTB | CITED4 |
| PRDX1 | TUBB | PTPRZ1 | AL117329.1 | ASB2 | KRT19 | DSC2 | EMP3 |
| TOMM5 | UBE2C | RBBP8 | CCDC91 | B4GALT5 | LGALS1 | DSG3 | FGFBP1 |
| TUBA1B | CENPF | TRPS1 | DLG1 | CCND1 | MIR205HG | DSP | HMGA2 |
| EIF5A | H2AFZ | ADGRL3 | EXOC6B | CTNND1 | NDRG1 | FABP5 | IGFBP6 |
| FGFBP1 | MKI67 | AL117329.1 | FRMD6 | CTSC | NDUFA4L2 | HSPB1 | IL1B |
| HSP90AA1 | TPX2 | ARHGAP24 | IMMP2L | EIF4EBP1 | PDLIM1 | S100A8 | INHBA |
| RANBP1 | TUBB4B | CACNB4 | INPP4B | ELF3 | PGK1 | SERPINB1 | ITGB4 |
| RPS26 | UBE2S | CADPS2 | KIAA1217 | FABP5 | TPM2 | SLPI | KRT14 |
| SNRPF | ANLN | CDK6 | PATJ | GCHFR | ADIRF | SPRR1B | KRT19 |
| SRM | HMGB1 | COL8A1 | PTPRK | GLIS3 | ANGPTL4 | CA2 | KRT6A |
| ATP5MC3 | KIF20B | CPA6 | SBF2 | GUCY1A1 | BST2 | CD24 | LAMA3 |
| DBI | KPNA2 | DST | TRPS1 | HES4 | BTG1 | CDA | LAMB3 |
| DCTPP1 | PTTG1 | FGFR1 | AC012494.1 | KRT8 | CKB | CENPW | LAMC2 |
| FDPS | SMC4 | FOXP1 | ANK3 | MARCKS | CTSD | CLTB | NDRG1 |
| HSPD1 | ARL6IP1 | FTX | AP000676.5 | MT1X | EPB41L4A-AS1 | CRABP2 | PLAU |
| NCL | ASPM | GMDS | CCDC18-AS1 | MTSS1 | FAM3C | CSTA | RBP1 |
| NDUFA4 | CENPW | KCNQ5 | DLEU1 | PLD1 | FTL | CTSC | S100A10 |
| PDCD5 | PCLAF | LAMA1 | EHBP1 | RDH10 | GAS5 | JUP | S100A11 |
| POLR2L | TUBA1C | LPCAT2 | ESYT2 | S100A9 | HCFC1R1 | KRT13 | S100A14 |
| PSMB3 | CALM2 | LRMDA | FMNL2 | SHANK2 | IFI6 | KRT14 | S100A16 |
| SLC3A2 | CCNB1 | MALAT1 | KANK1 | SULF2 | IFITM3 | KRT19 | SFN |
| SNRPB | CCNB2 | MAPK10 | KDM4C | TF | IGFBP2 | KRT23 |  |
| SNRPG | CDC20 | PAM | LPP | TMSB4X | IGFBP6 | KRT5 |  |
| TXN | CENPE | PDZRN4 | MAML3 |  | JUNB | LYPD3 |  |
|  | DEK | PRELID2 | MAPK10 |  | LDHA | MMP7 |  |
|  | DLGAP5 | PSD3 | MED13L |  | LEMD1 | NDRG1 |  |
|  | DUT | PTPN14 | MIR31HG |  | MT1X | PI3 |  |
|  | H2AFV | SEMA3C | MTRNR2L12 |  | NUPR1 | PKP1 |  |
|  | HMGN2 | TRIM2 | NHSL1 |  | P4HA1 | RAB11FIP1 |  |
|  | HMMR |  | NRIP1 |  | PFDN5 | RHOV |  |
|  | MAD2L1 |  | PARD3 |  | PHLDA1 | S100A10 |  |
|  | PRC1 |  | PCDH7 |  | PNRC1 | S100A11 |  |
|  | RRM2 |  | PDE4D |  | PTHLH | S100A6 |  |
|  | TK1 |  | PTPRD |  | RPL10 | S100A7 |  |
|  | UBE2T |  | RIMS2 |  | RPS27 | SAT1 |  |
|  |  |  | SCHLAP1 |  | SERPINA3 | SERPINB4 |  |
|  |  |  | SHANK2 |  | SNHG7 | SFN |  |
|  |  |  | SLC22A23 |  | SOX4 | SPINK5 |  |
|  |  |  | SPIDR |  | TINAGL1 | SPRR2D |  |
|  |  |  | STAG1 |  | TMEM45A |  |  |
|  |  |  | TENM2 |  | TPI1 |  |  |
|  |  |  | TMTC2 |  | TUBA1A |  |  |
|  |  |  | TP63 |  | TXNIP |  |  |
|  |  |  | TRIO |  |  |  |  |
|  |  |  | WSB1 |  |  |  |  |
|  |  |  | ZBTB20 |  |  |  |  |
|  |  |  | ZFAND3 |  |  |  |  |

ITH: intratumor heterogeneity

**Supplementary Table 6.** AREG mRNA expression in human HNSCC cell lines (CCLE data)

| Cell lines | AREG (log_2_FPKM) |
| --- | --- |
| HSC4 | 3.280956 |
| PECAPJ34CLONEC12 | 6.367371 |
| SCC15 | 8.685029 |
| SCC4 | 8.63492 |
| UPCISCC026 | 6.335926 |
| UPCISCC029A | 1.244887 |
| UPCISCC040 | 6.224581 |
| UPCISCC074 | 6.65263 |
| UPCISCC152 | 3.054848 |
| SNU1041 | 7.415488 |
| SNU1066 | 5.867155 |
| SNU1076 | 4.219556 |
| SNU1214 | 6.231317 |
| BICR22 | 6.684398 |
| BICR31 | 6.981853 |
| BICR78 | 5.848998 |
| CAL33 | 5.808128 |
| SNU46 | 3.522307 |
| BICR6 | 4.703211 |
| BICR16 | 7.392575 |
| BICR56 | 7.44427 |
| YD8 | 1.214125 |
| YD10B | 6.901712 |
| YD15 | 5.657926 |
| YD38 | 9.076174 |
| PECAPJ49 | 8.109883 |
| H103 | 9.211572 |
| H357 | 7.800771 |
| H376 | 8.571753 |
| H413 | 4.996841 |
| UPCISCC131 | 4.697663 |
| UPCISCC154 | 4.782409 |
| UPCISCC200 | 4.556429 |
| H157 | 4.163499 |
| SNU899 | 5.91576 |
| PECAPJ15 | 6.756356 |
| PECAPJ41CLONED2 | 7.44791 |
| BHY | 7.944507 |
| UPCISCC116 | 5.523248 |
| UPCISCC111 | 2.801159 |
| DETROIT562 | 4.984134 |
| RPMI2650 | 0.042644 |
| SCC25 | 5.393691 |
| SCC9 | 8.712493 |
| CAL27 | 3.989139 |
| SW579 | 0.545968 |
| A253 | 5.500802 |
| FADU | 5.302319 |
| KOSC2 | 3.291309 |
| KON | 2.526069 |
| OSC20 | 7.00304 |
| OSC19 | 8.035734 |
| HSC2 | 5.511595 |
| HSC3 | 10.03604 |
| SAT | 5.074677 |
| HSQ89 | 1.356144 |
| T3M5 | 8.76454 |
| SAS | 7.614342 |
| CA922 | 5.313246 |
| HO1U1 | 4.050502 |

**Supplementary Table 7**

| qPCR primer | Forward (5' to 3') | Reverse (5' to 3') |
| --- | --- | --- |
| TBP | TGTATCCACAGTGAATCTTGGTTG | GGTTCGTGGCTCTCTTATCCTC |
| CAV1 | CCAAGGAGATCGACCTGGTCAA | GCCGTCAAAACTGTGTGTCCCT |
| SFN | TGCTGGACAGCCACCTCATCAA | GGCTGAGTCAATGATGCGCTTC |
| ITGB4 | AGGATGACGACGAGAAGCAGCT | ACCGAGAACTCAGGCTGCTCAA |
| KRT6A | GAGGAGATTGCTCAGAGAAGCC | CAATCTCCTGCTTGGTGTTGCG |
| LAMA3 | TAGAGGAAGCCTCTGACACAGG | CCGATAGTATCCAGGGCTACAAC |
| LAMB3 | GTCACAGAGCAGGAGGTGGCT | GCTTCTGTCAAGACTCTCCAGG |
| LAMC2 | TACAGAGCTGGAAGGCAGGATG | GTTCTCTTGGCTCCTCACCTTG |
